# MS1Connect: a mass spectrometry run similarity measure

**DOI:** 10.1101/2022.01.12.476125

**Authors:** Andy Lin, Brooke L. Deatherage Kaiser, Janine R. Hutchison, Jeffrey A. Bilmes, William Stafford Noble

## Abstract

Interpretation of newly acquired mass spectrometry data can be improved by identifying, from an online repository, previous mass spectrometry runs that resemble the new data. However, this retrieval task requires computing the similarity between an arbitrary pair of mass spectrometry runs. This is particularly challenging for runs acquired using different experimental protocols. We propose a method, MS1Connect, that calculates the similarity between a pair of runs by examining only the intact peptide (MS1) scans, and we show evidence that the MS1Connect score is accurate. Specifically, we show that MS1Connect outperforms several baseline methods on the task of predicting the species from which a given proteomics sample originated. In addition, we show that MS1Connect scores are highly correlated with similarities computed from fragment (MS2) scans, even though this data is not used by MS1Connect. The MS1Connect software will be made available upon acceptance at https://github.com/bmx8177/MS1Connect.

## 1 Introduction

Over the years a wealth of proteomics data has been deposited into online repositories such as PRIDE^1^ and MassIVE.^2^ Researchers may be interested in finding data in these online repositories that are similar to their own data in order to perform some joint analysis. However, it is difficult to identify which repository runs should be analyzed with your data. This is because it is hard to measure the similarity of a pair of runs, especially across different studies. Specifically, biologically irrelevant differences in sample preparation, liquid chromatography, and instrument parameters all affect the resulting data.

Currently, few methods exist for measuring the similarity of a pair of proteomics runs. One metric is to count the number of confidently detected peptides in common.^3^ Unfortunately, this method requires knowing the species composition of the samples, which is not always known, and requires a database search. Another previously developed method for measuring the similarity between a pair of mass spectrometry runs directly compares the set of peptide fragment spectra (MS2 scans) from each run. Conceptually this method works by calculating the spectra dot product for all pairs of spectra in two runs and then counting the fraction of scores above a threshold.^4^ This approach has been successfully used for species identification,^5^ molecular phylogenetics,^6^ and differentiating between experimental protocols.^7^

Unfortunately these MS2 based methods do not account for differences in how MS2 scans are collected. MS2 spectra collected by data-independent acquisition (DIA), data-dependent acquisition (DDA), and targeted analyses are all different from each other. Therefore, new methods are needed that can measure the similarity between a pair of proteomics runs regardless of how the data was acquired.

In this work, we describe a new method, MS1Connect, that only uses intact peptide scans (MS1 scans) to calculate the similarity between a pair of runs. Since MS1 scans are always collected in the same way, this data can be used to compare targeted, DDA, and DIA data. To our knowledge, MS1 data has not been used to measure the overall similarity between a pair of proteomics runs. However, several methods have been developed to align MS1 features maps of a pair of proteomics runs,^8–10^ but these methods do not report an overall similarity score.

Our method, MS1Connect, frames scoring the similarity between a pair of proteomics runs as a maximum bipartite matching problem. A bipartite graph consists of two disjoint sets of vertices and a set of edges that connect the two sets of vertices. In a maximum bipartite matching problem, the goal is to select a subset of edges in a bipartite graph that maximizes some objective value subject to some constraint. In our setting, each of the two disjoint sets of vertices represent the set of MS1 features found in a run, and edges link MS1 features, in different runs, whose masses match within some tolerance. In addition, we require that every MS1 feature be associated with at most one edge in the set of selected edges.

The MS1Connect objective function consists of a weighted combination of three modular terms and a fourth supermodular term. Modular and supermodular functions are both types of set functions. In a modular function the sum is equals to the sum of its parts. More specifically, given two sets of disjoint items *X* and *Y* and a scoring function *f, f* (*X*) + *f* (*Y*) = *f* (*X* ∪ *Y*). On the other hand, for a supermodular function, the sum is greater than the sum of its parts. In this case *f* (*X*) + *f* (*Y*) ≤ *f* (*X* ∪ *Y*). While there exists a robust set of literature dedicated to the theory of supermodular maximization,^11–14^ these functions have rarely been applied to the analysis of biological data. The one example we are aware of uses a surrogate function to estimate the supermodular relationship between fragment ions in database searching.^15^

Each of the MS1Connect terms measures a different aspect of proteomics run similarity. The first modular term (*M*_1_) favors solutions with a large number of edges. The second modular term (*M*_2_) favors selection of edges with high intensity values. The third modular term (*M*_3_) favors edges with small normalized retention time shifts. We define the normalized retention time shift as the difference in normalized retention times between the two MS1 features in an edge. Finally, the fourth supermodular term (*M*_4_) favors solutions with pairs of edges that are similar to each other. Two edges are similar if they have similar normalized retention time shifts and if the normalized retention times of the MS1 features, in the same run, are similar to each other.

We show evidence that the MS1Connect score accurately measures the similarity between two proteomics runs. There are many ways to define whether two runs are similar to each other. For this work we focus on the task of species prediction. We show that MS1Connect scores outperforms baselines for predicting the species a sample originated from. In addition, we show that MS1Connect scores are able to recapitulate similarities based off MS2 spectra. Specifically, we show a high correlation between MS1Connect scores and the Jaccard index between the sets of confidently detected peptides for a pair of runs.

## 2 Approach

### 2.1 Representation of a mass spectrometry run

We represented each tandem mass spectrometry run as a bag of MS1 features, where each feature nominally corresponds to a peptide detected in a set of precursor scans. Each MS1 feature is represented as a tuple of four values: *m/z*, intensity, charge, and retention time (in seconds). All four of these values are reported by pyOpenMS,^16^ the tool we use for MS1 feature detection. To generate the input files for pyOpenMS, we used Proteowizard version 3.0^17^ to convert Thermo RAW files to .mzML format.^18^ We use the *N* most intense MS1 features to represent each run, where *N* is a hyperparameter. In Supplemental Section 1.1 we discuss how we normalized the values returned by pyOpenMS.

### 2.2 Mass spectrometry run matching as a maximum bipartite matching problem

We frame the measurement of the similarity between a pair of mass spectrometry runs as a maximum bipartite matching problem. In this approach, we aim to select a set of edges in a given bipartite graph that achieves a maximum objective value.

A bipartite graph *G* = (*U, V, E*) consists of two disjoint sets of vertices, *U* and *V*, and a set of edges 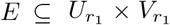, where each edge *e* connects a vertex *u* ∈ *U* to a vertex *v* ∈ *V*. For our specific formulation, *U* and *V* are the sets of MS1 features from two different mass spectrometry runs, *r*_1_ and *r*_2_, and the edges *E* link MS1 features between *U* and *V* (Supplemental Figure 1). For this reason, we have one bipartite graph 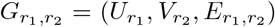 associated with every run pair *r*_1_, *r*_2_, where 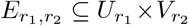. For notational simplicity, we drop the *r*_1_, *r*_2_ subscripts except when they are needed for run-pair disambiguation.

We include in the graph edges between all pairs of MS1 features whose charges match and whose *m/z* values match within some tolerance *δ*_1_ (in units of ppm):

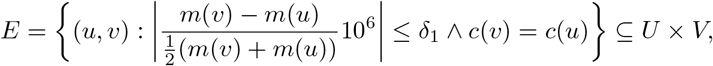

where *m*(*u*) is the *m/z* of *u, m*(*v*) is the *m/z* of *v, c*(*v*) is the charge of MS1 feature *v*, and *c*(*u*) is the charge of *u*. By connecting MS1 features with the same charge and similar *m/z*, these edges attempt to connect the same peptide precursor.

The goal in maximum bipartite matching is to select a set of edges *A* that achieves a maximum score, as measured by a specified objective function *S*, subject to some matching constraints. Specifically, because we expect that each peptide precursor will be detected at most once per run, we require that a valid matching connects each feature in *U* to at most one feature in *V* and vice versa. More formally, for two runs *r*_1_ and *r*_2_, we want to choose a subset of edges *A* ⊆ *E* that achieves a maximum value of the objective defined below subject to the following constraints: ∀*e ∈ A*, degree(*u*(*e*)) ≤ 1 and degree(*v*(*e*)) ≤ 1, where *u*(*e*) retrieves the relevant MS1 feature from *U* and *v*(*e*) retrieves the corresponding MS1 feature in *V*. To designate these constraints, we define ℰ_*U*_ = {*A* ⊆ *E* : ∀*e* ∈ *A*, degree(*u*(*e*)) ≤ 1} which is the set of all subsets of edges that satisfy by the required degree constraint of the corresponding nodes on the *U* side, and we correspondingly define ℰ_*V*_ = {*A* ⊆ *E* : ∀*e ∈ A*, degree(*v*(*e*)) ≤ 1} for the *V* side.

### 2.3 Scoring a candidate matching

Our handcrafted score function 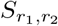 uses a maximum matching approach which maximizes an objective that consists of a weighted combination of four terms. The score for two runs *r*_1_, *r*_2_ is defined as follows:

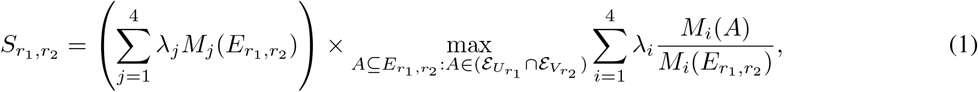

where *M*_1_ through *M*_4_ are terms that are defined below, *λ*_1_ through *λ*_4_ are convex mixture hyperparameters that weight the relative importance of each term, and 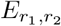 is the set of valid edges between runs *r*_1_ and *r*_2_. Each term in the inner maximization is normalized by the score for the full set *E* with no matching constraints. The inner maximization maximizes over all subsets of edges that satisfy by both degree constraints, and that is indicated via 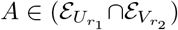 since if *A* is a member of both 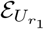 and 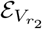, then no node incident to any edge has more than degree one in the matching. The leading summation serves to un-normalize the objective value via a multiplicative (*r*_1_, *r*_2_)-dependent constant. This two-step normalization and un-normalization process is used to make the *λ* hyperparameters interpretable (i.e., each *λ*_*i*_ indicates the relative contribution of each term) and also to ensure that the scores are calibrated over multiple distinct pairs of runs. As a result of the normalization process, the four *λ* values must sum to one and range between zero and one, inclusive. We next describe the *M*_*i*_ terms, each of which measures a different aspect of proteomics run similarity (Figure 1).

**Figure 1:**
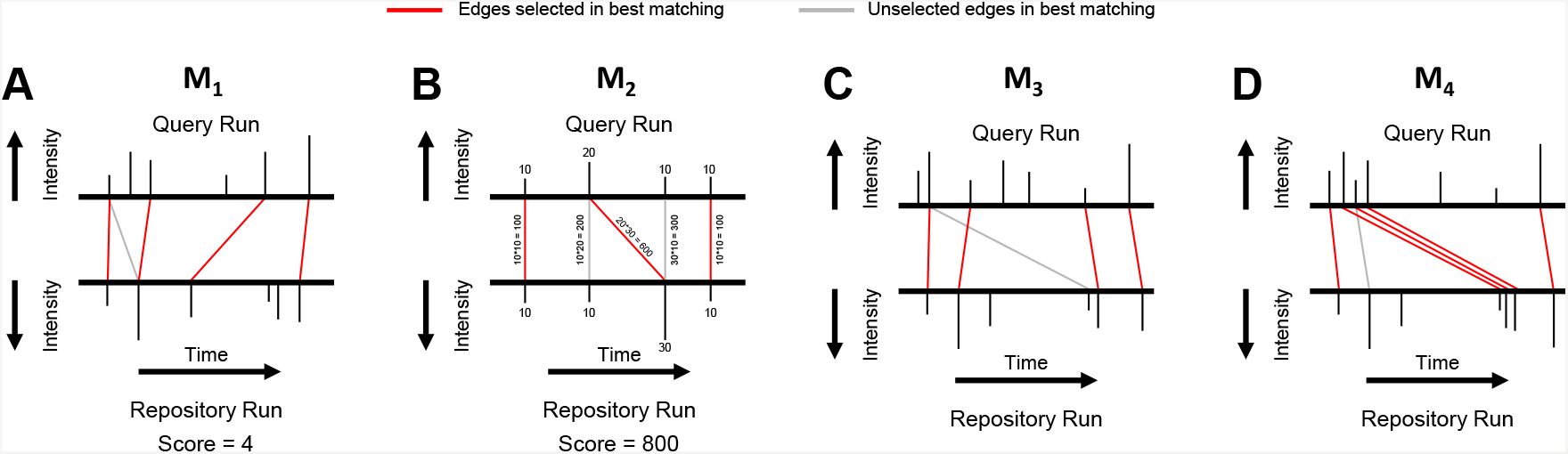
Schematic of four MS1Connect terms. This figure illustrates what each term of the MS1Connect objective function focuses on when scoring a matching. Each subplot shows bipartite graph formed by a pair of mass spectrometry runs. The red and gray lines are edges that link MS1 features with similar *m/z*. Note that the *m/z* axis is not shown. In addition, note that, in a specific matching, each MS1 feature is associated with at most one edge. (A) The *M*_1_ term favors solutions that contain a large number of edges. (B) The *M*_2_ term favors solutions that select edges with high intensity values. (C) The *M*_3_ term favors edges that have small normalized retention time shifts. (D) The *M*_4_ term favors solutions that contain many similar pairs of edges. The best solution chooses the three red edges in the middle instead of the gray edge because these three edges are similar to each other. This similarity indicates that the large retention time difference is correct and due to systematic differences in chromatography conditions.

The first term *M*_1_ counts the number of edges selected in a solution: *M*_1_(*A*) = |*A*|. Thus, this term favors solutions that contain a large number of edges. Runs that are similar will share many of the same peptides. As a result, these peptides will be linked by edges, thereby generating many edges.

The second term sums the product of the two intensities associated with each edge found in a selected matching, *M*_2_(*A*) = Σ_*e*∈*A*_ *I*(*u*(*e*))*I*(*v*(*e*)), where *I*(*u*) in the intensity of feature *u* and *I*(*v*) in the intensity of feature *v*. This term favors solutions that select edges with high intensity values. The intensity of an MS1 feature produced by a peptide is a function of the sequence and abundance of that peptide. Runs that are similar to each other should have similar peptide expression levels. By multiplying the two intensities together, we focus on the most intense peptide features.

The third term sums the negative exponent of the absolute difference of the normalized retention time shift of all the selected edges: *M*_3_(*A*) = Σ_*e*∈*A*_ exp(−*α*|*s*(*e*)|), where *α* is a hyperparameter that determines how quickly the exponential function decreases and *s*(*e*) is the normalized retention time shift between *u* and *v*: *s*(*e*) = *t*(*v*(*e*)) − *t*(*u*(*e*)), where *t*(*u*) and *t*(*v*) are the normalized retention times of MS1 features *u* and *v*, respectively. This term favors edges that have small normalized retention time shifts. Such edges are better because they are more likely to link the same peptide feature. This is because we expect the time it takes for peptides to elute off the column, in similar chromatography conditions, to be similar from one run to the next.

Finally, the fourth term scores pairs of edges rather than a single edge. This term favors solutions that contain many similar pairs of edges. We say a pair of edges are similar to each other when they have similar normalized retention time shifts and the normalized retention times of the MS1 features, in the same run, are similar to each other. We define the term as

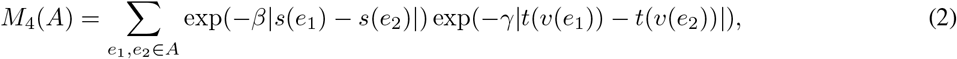

where *β* and *γ* are hyperparameters. The first exponent is large when the normalized retention time shifts of the two edges are similar. On the other hand, the second exponent is large when pairs of MS1 features in the same run have similar normalized retention times. Altogether, this term scores edge pair similarity while allowing for systematic retention time shifts between two different mass spectrometry runs. Such shifts are common, reflecting differences in chromatography conditions between runs. Note that, for efficiency, we set *M*_4_ to zero when |*s*(*e*_1_) − *s*(*e*_2_)| *>* 0.01 or when |*t*(*v*(*e*_1_)) − *t*(*v*(*e*_2_))| *>* 0.01.

### 2.4 Selecting the best matching

In order to choose the best matching, we must choose a feasible subset of edges that maximizes our objective function, that is, we must compute

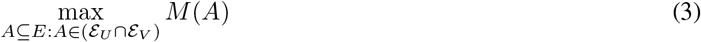

where 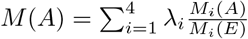. The first thing to note is that *M* (*A*) is the summation over entries of a |*U* ||*V* |×|*U* ||*V* | matrix. That is, 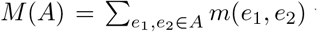 where 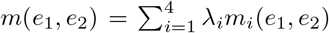 and where *m*_*i*_(*e*_1_, *e*_2_) is the element of the corresponding sub-objective *M*_*i*_ defined above (that is, 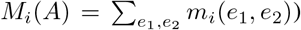. Note that for *i* ∈ {1, 2, 3} we have that *m*_*i*_(*e*_1_, *e*_2_) = 0 whenever *e*_1_ ≠ *e*_2_. We also note that *m*_4_(*e*_1_, *e*_2_) = 1 whenever *e*_1_ = *e*_2_. Hence, we can define *m*(*e*_1_, *e*_2_) as follows:

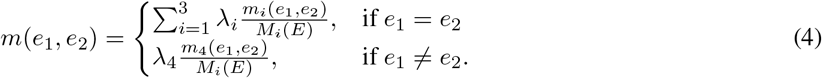

Ordinarily, computing such a maximization over an exponential number of subsets would be intractable. It turns out, however, that this maximization problem exhibits useful structure that allows an efficient algorithm to be used to obtain an approximate solution. Specifically we use the greedy algorithm to solve an instance of supermodular maximization subject to matroid constraints. In Supplemental Section 1.2 we discuss our approach to select the best matching as well as the approximation bound of the solution provided by the greedy algorithm.^11–13,19,20^

Lastly, we note again that the above is described for a generic bipartite graph *G* = (*U, V, E*). In practice, we have one distinct bipartite graph for each pair *r*_1_, *r*_2_ of runs; hence, the greedy algorithm is run for each run pair.

## 3 Methods

Additional methods can be found in the Supplement.^21–28^

### 3.1 MS1Connect versions

Since the objective function of MS1Connect contains four terms, we compared five different versions of MS1Connect against each other. Four of the versions used a single term of the objective function while the fifth version used all four terms of the objective function. We notate the version of MS1Connect that only uses the *M*_1_ term as “MS1Connect (*M*_1_ only)”, and we notate the version of MS1Connect that uses all four terms as “MS1Connect (*M*_1_ – *M*_4_)”.

### 3.2 Baseline similarity measures

In order to ensure that our objective obtained sensible results, we compared MS1Connect against several baseline similarity measures. In general, our baseline similarity measures are based on binning the MS1 features and computing scalar products. More specifically, we represent the two runs, *X* and *Y*, as two matrices with dimensions *m* and *n*, where *m* is the number of *m/z* bins and *n* is the number of retention time bins. The value in each cell is the number of MS1 features whose discretized *m/z* and retention time fall into that bin. Both *δ*_2_, the *m/z* bin width, and *n*, the number of retention time bins, are hyperparameters.

Our baseline scores are defined as

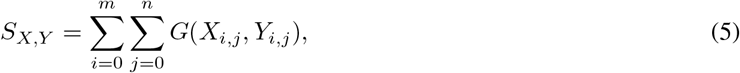

where *X* and *Y* are the matrix representations of the proteomics runs and *G* is the product, maximum, or minimum of the two input values. Alternatively, *G* could be defined as *G*(*S, T*) = 𝟙_{*S*≥1}_𝟙_{*T* ≥1}_, where 𝟙 is the indicator function. Finally, we considered the baselines that do not consider retention time (*n* = 1) separately from baselines that do consider retention time (*n* ≥ 2) for a total of eight different baselines.

### 3.3 Evaluation metrics

We use two different measures to quantify the performance of a given MS1 similarity score at predicting the metadata label of a query run given a repository of runs with known metadata labels.

The first performance measure is the per-query average precision (QAP), defined as the average precision across all queries *q* ∈ *R*:

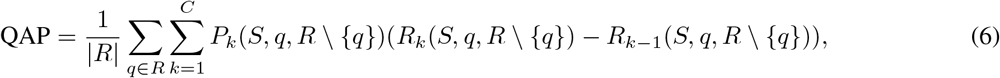

where *P*_*k*_(*S, q, R*) and *R*_*k*_(*S, q, R*) are the precision and recall, respectively, after *k* repository runs have been retrieved from *R* using query *q* and similarity *S*. Note that the query run *q* is not included in the ranking when computing the precision and recall.

The second measure, aggregate average precision (AAP), is similar to QAP except that the average precision is calculated once on an aggregated list of similarity scores. This aggregated list is produced by sorting together all pairs of runs, considering only the upper triangle of the run-by-run matrix. In this work we focused on the QAP because these two performance metrics are highly correlated (Supplemental Figure 2).

## 4 Results

### 4.1 MS1Connect scores can be used for species prediction

To determine whether the MS1Connect score successfully measures the similarity of a pair of runs, we investigated whether these scores can be used to predict the species label of a proteomics run. Given a set of runs, we designated a single run as the query run and the remaining runs as the repository. The repository runs were then ordered by their similarity to the query run. Then, the average precision was calculated based on whether the repository runs had the same species label as the query run. We repeated this process for each of the runs in the dataset to calculate our two metrics, QAP and AAP. MS1Connect has nine different hyperparameters (Table 1), and to determine which set of hyperparameters achieved the best performance on the training set, as measured by QAP, we sampled the hyperparameter space using a random grid search.

**Table 1:** QAP and AAP. A table of the per-query average precision (QAP) and the aggregate average precision (AAP) of the best performing hyperparameter set for each method in the species training and test datasets. Bold values denote the best and second best performance for each column. MS1Connect (*M*_1_ – *M*_4_) and MS1Connect (*M*_4_ only) had the overall best performance. Note that the two supermodular methods outperformed all other methods.

| method | species training set |  | species test set |  |
| --- | --- | --- | --- | --- |
|  | QAP | AAP | QAP | AAP |
| MS1Connect ($M_1 - M_4$) | **0.8115** | **0.7301** | **0.8483** | **0.7983** |
| MS1Connect ($M_1$ only) | 0.7873 | 0.6992 | 0.8112 | 0.7672 |
| MS1Connect ($M_2$ only) | 0.7096 | 0.5801 | 0.6686 | 0.5806 |
| MS1Connect ($M_3$ only) | 0.7891 | 0.7000 | 0.8130 | 0.7675 |
| MS1Connect ($M_4$ only) | **0.8076** | **0.7322** | **0.8497** | **0.7993** |
| # $m/z$ bins in common (indicator) | 0.7847 | 0.6978 | 0.8211 | 0.7855 |
| # $m/z$ + RT bins in common (indicator) | 0.7677 | 0.6856 | 0.8083 | 0.7509 |
| # $m/z$ bins in common (product) | 0.7771 | 0.7062 | 0.7951 | 0.7529 |
| # $m/z$ + RT bins in common (product) | 0.7591 | 0.6838 | 0.8107 | 0.7532 |
| # $m/z$ bins in common (min) | 0.7831 | 0.6979 | 0.8225 | 0.7868 |
| # $m/z$ + RT bins in common (min) | 0.7665 | 0.6857 | 0.8092 | 0.7514 |

We found that MS1Connect scores can be successfully used for predicting what species a sample was generated from (Table 1). Comparing the various versions of MS1Connect against each other, we discovered that the two supermodular versions of MS1Connect, MS1Connect (*M*_1_ – *M*_4_) and MS1Connect (*M*_4_ only), had the best performance. Specifically, these two versions had the best or second best performance, as measured by QAP or AAP, in both the training and test data. These two versions are supermodular because they incorporate the supermodular *M*_4_ term. The higher performance of these two supermodular versions of MS1Connect, compared to the three modular versions of MS1Connect, suggests that using supermodularity may lead to higher performance. In Supplemental Note 3.1 we discuss a possible reason why supermodular methods outperform modular methods.

Next, we compared the performance of MS1Connect against our baseline similarity methods (Table 1). We found that the supermodular MS1Connect methods outperformed all of the baselines. In addition, we found the performance of MS1Connect (*M*_1_ only) and MS1Connect (*M*_3_ only) to be similar to the performance of the baselines while MS1Connect (*M*_2_ only) under-performed the baselines. Note that Table 1 does not include the performance of the two baselines where *G* is the maximum of the two input values as these methods exhibited poor performance (≤ 0.13 for both QAP and AAP in the training set).

After we compared MS1Connect against the baselines, we compared the various baselines against each other. We found that performance generally decreased when retention time was considered (Table 1). This result may seem unexpected because retention times, in principle, should improve the specificity of a matching. However, binning retention times could lead to a significant number of edge effects because of systemic retention time shifts that reflect differences in chromatography conditions. As an example, the number of MS1 features in common between two *C. albicans* runs, 1512006-2-TRIS1-10-Calbicans.raw and Control-60min R3.raw, decreased by 30.8% after increasing the number of retention time bins from one to two.

Following the quantitative comparison, we examined the MS1Connect scores in the species training data to understand whether these scores made qualitative sense. We expected that runs generated by the same experiment should be highly similar to each other. Beyond that, we also expected runs from the same species to also be similar with each other.

Using the hyperparameter set that yielded the best performance, as measured by QAP on the training data, we visualized the MS1Connect scores between all pairwise runs in the species training dataset as a heatmap (Figure 2). In this heatmap the rows and columns are ordered by expected similarity. In general, the resulting heatmap matched our expectation. For example, the strong diagonal component in the heatmap resulted from runs that were generated by the same experiment. In addition, the block-diagonal structure (delimited in the figure by the solid white lines) showed that runs generated from the same species tend to have high scores. For example, all the runs generated by *S. aureus* and *C. albicans* generally only have high scores with each other. On the other hand, we also found a few examples of runs whose similarity profile did not match our expectation. One prominent example is a set of five *E. coli* runs from the same experiment that only had high MS1Connect scores when comparing the same run to itself. A second example is a set of five *A. thaliana* runs from the same experiment where a run often did not have a high MS1Connect score to itself. This result occurred because these 10 runs have few MS1 features. In general, larger MS1Connect scores can be achieved when a run has a larger number of MS1 features. In this specific situation, these 10 runs have the fewest number of MS1 features in the overall dataset.

**Figure 2:**
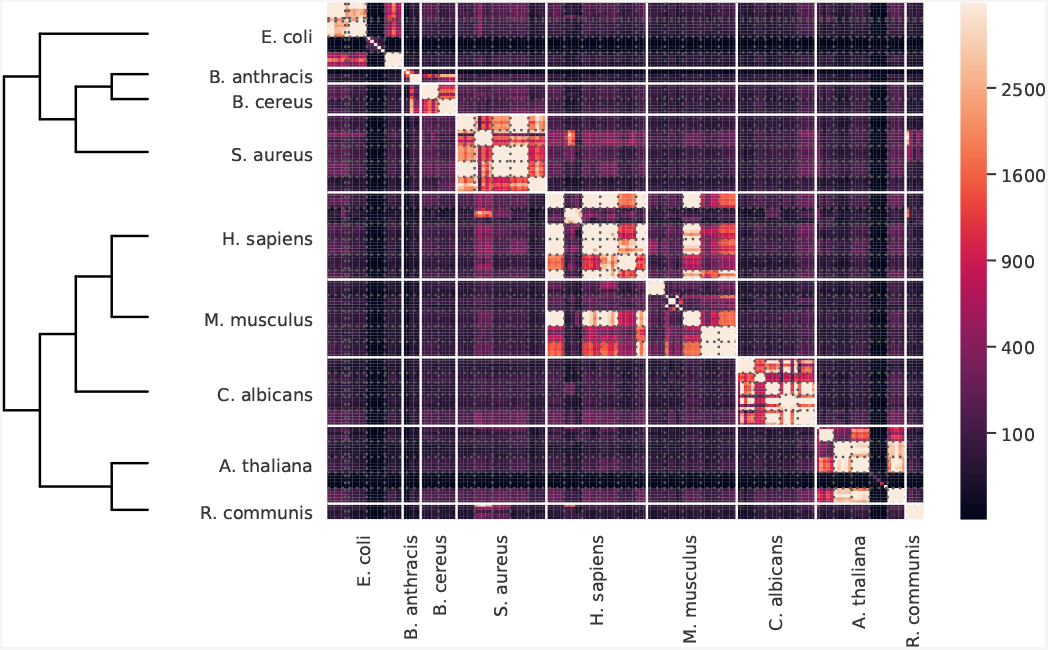
MS1connect for species prediction. A heatmap and dendrogram of MS1Connect scores showing the structure of our species training dataset. Each cell is colored by the MS1Connect score between a pair of runs. The solid white lines denote the border between different species while the dotted gray lines delineate the border between different experiments (PRIDE ID). The denodrogram shown is the phylogenetic tree of the nine species found in the training dataset.

One unexpected finding was that MS1Connect scores may be sensitive enough to detect inter-species relationships. Considering the human and mouse runs, the MS1Connect scores indicated that these two species have some degree of similarity. This finding fits with our phylogenetic understanding of these two species, as human and mice are both mammals. These two species are highly similar to each other in the context of our dataset, which also included bacteria, fungi, and plants. Another example is that *B. anthracis* and *B. cereus* runs are similar to each other, which is expected since these two species are in the same genus. In the species test data, we saw comparable results with the two gram-negative bacterial species, *E. coli* and *S. enterica*, being similar to each other (Supplementary Figure 5).

A close inspection of the heatmap showed an unexpectedly high similarity between a group of *S. aureus* runs and a group of *H. sapiens* runs (Figure 2). The *S. aureus* runs originated from a study that used both cultures and clinical samples of *S. aureus*. We hypothesized that these runs originated from clinical samples and therefore contained a large number of human peptides. This type of contamination would cause these bacterial runs to be similar to human runs. We were unable to confirm this information from the PRIDE metadata, because the submission did not track which samples were cultured-based and which samples were clinically-based. However, a database search of the *S. aureus* runs against a concatenated *S. aureus* and *H. sapiens* proteome detected a large number of human peptides, supporting our hypothesis.

### 4.2 M1Connect scores replicate database search results

Our analyses so far suggest that MS1Connect scores can successfully measure the similarity between pairs of proteomics runs. Next, we compared whether our MS1-based similarity method replicates similarities generated from MS2 data. We expect that these two methods should generally agree with each other.

To test this hypothesis, we compared the MS1Connect scores against the similarities generated by a MS2-based method on a set of samples with known species composition. For the MS2-based method, we calculated the Jaccard index between the sets of confidently detected peptides at a 1% FDR threshold for a pair of runs. The dataset we used contained a set of 16 samples, labeled “A” through “P”, where each sample contained one, two, or three different organisms (Supplementary Table 6). Each sample, except for the sample labeled “P” was run on a mass spectrometer four times. Three of the runs were analyzed by a single instrument in one laboratory while the fourth run was analyzed on a different instrument in a second laboratory. The two laboratories are independent from each other, and therefore the samples were run with differing instrument parameters and chromatography conditions.

Our results showed that MS1Connect scores indeed have a high correspondence to similarities measured from MS2 data. A heatmap where the upper triangle is MS1Connect scores and the lower triangle is the Jaccard index showed that these two methods are highly consistent with each other (Figure 3). In addition, we found that these two scores are highly correlated, with a Spearman rank correlation of 0.91. Overall, this result showed that MS1Connect scores can replicate MS2-based analyses. In addition, our result suggested that MS1Connect scores can measure the similarity between runs that contain multiple species in the face of differing chromatography conditions and instrumentation.

**Figure 3:**
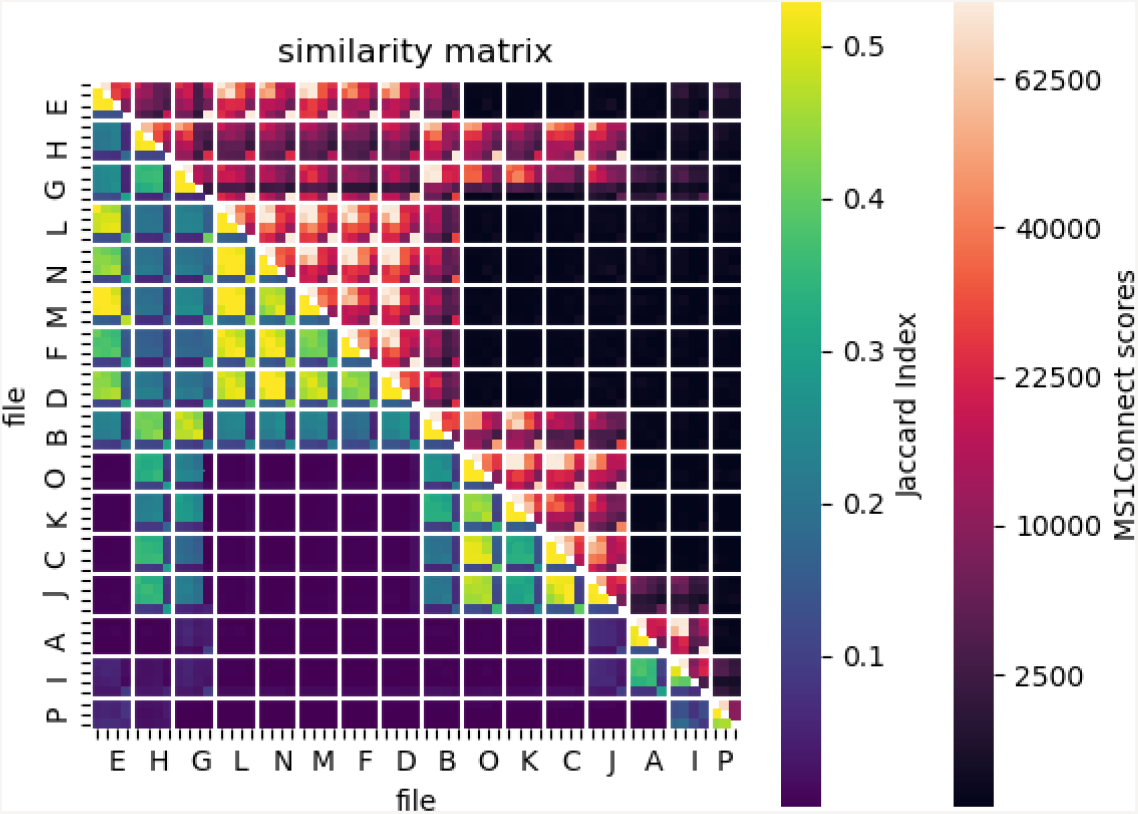
Similarity between MS1Connect scores and MS2-based Jaccard index. A heatmap of similarities scores where the upper triangle is MS1Connect scores and the lower triangle is the Jaccard index between the two sets of confidently detected peptides at a 1% FDR threshold for the multi-species dataset. These two methods are highly correlated with a Spearman rank correlation of 0.91. The solid white lines denote the border between samples. A description of what species are in each sample can be found in Supplementary Table 6. Within a box delineated by the white lines, the first three rows are the runs from one laboratory while the fourth row is the run from a second laboratory.

A distinct checkerboard pattern in the lower triangle of the heatmap occurred because three out of the four runs from each sample are run on a single instrument while the fourth is run on a second instrument (Figure 3). This result matched our expectation since the three runs from the same instrument should be more similar to each other than the run from the second instrument. This pattern, while not as prominent, also exists on the MS1Connect side of the heatmap and further showed that MS1Connect scores can replicate database search results.

### 4.3 MS1Connect scores can identify mislabeled runs

Having shown the ability of MS1Connect to accurately measure the similarity between pairs of proteomics runs, we hypothesized that our method could be used to identify runs with mislabeled metadata. To this end, we analyzed data collected from seven different *Bacillus* species.^28^ Each species was analyzed by mass spectrometry six times. We calculated the MS1Connect scores between all of the runs. In addition, we calculated the Jaccard index between the sets of confidently detected peptides at 1% FDR threshold for all pairs of runs.

We found that MS1Connect scores identify runs whose metadata may have been mislabeled. Specifically, when visualizing a heatmap of the MS1Connect scores between all pairs of runs (upper triangle of Figure 4), we observed that one of the runs labeled as *B. tonyonesis* exhibits low similarity with other *B. tonyonesis* runs but has high similarity with other *B. cytotoxicus* runs (last row/column of the heatmap). A database search also confirmed this result (lower triangle of Figure 4). Together, these results strongly suggest that a single run labeled as *B. tonyonesis* should instead be labeled *B. cytotoxicus*. This mislabelling could have occurred in many ways. For example, during the sample preparation process a vial could have been mislabeled or there could have been an error during some pipetting step. Altnernatively, when placing samples into an autosampler, a vial could have been placed incorrectly onto the tray or the vial position could have been incorrectly recorded into the instrument.

**Figure 4:**
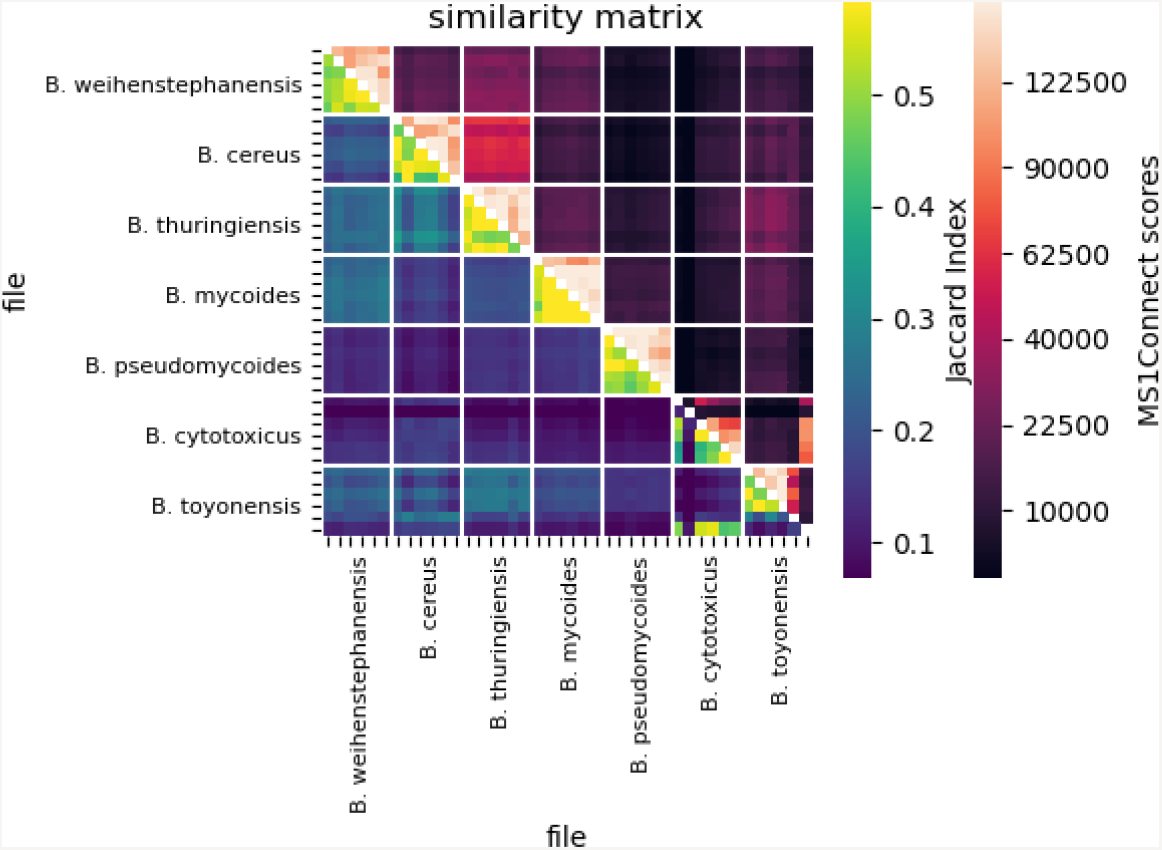
Possible identification of a mislabeled run. The figure is a heatmap of pairwise similarity scores, where the upper triangle is MS1Connect scores and the lower triangle is the Jaccard index between the two sets of confidently detected peptides at 1% FDR threshold for the *Bacillus* genus dataset. The x- and y-axis show the species labels of each run, as described by the PRIDE repository, with six runs per species. The solid white lines denote the border between species. Both the MS1Connect scores and the Jaccard index indicate that a run labeled as *B. tonyonesis* (last row/column of the matrix) was mislabeled and should be labeled *B. cytotoxicus*.

### 4.4 Analysis of hyperparameters

After the assessment of our method, we investigated the performance of MS1Connect as a function of the hyperparameters to determine if any of the hyperparameters were correlated with performance. Specifically, we studied the performance of the 2500 different hyperparameterizations of MS1Connect, as described in Supplemental Section 2.1, with respect to each hyperparameter.

This analysis revealed trends in the performance of various hyperparameterizations of MS1Connect as a function of *N* and *δ*_1_ (Figure 5). Specifically, we found that the best performance, as measured by QAP on the training dataset, occurred when the number of MS1 features used to generate the bipartite graph *N* was set to 4000 and when the *m/z* tolerance *δ*_1_ was set to 4 ppm. We note that 4 ppm is well within the range of precursor mass tolerances researchers typically use for database searching. On the other hand, we found no trends in the performance of MS1Connect as a function of the remaining seven hyperparameters (Supplementary Figures 6–7).

**Figure 5:**
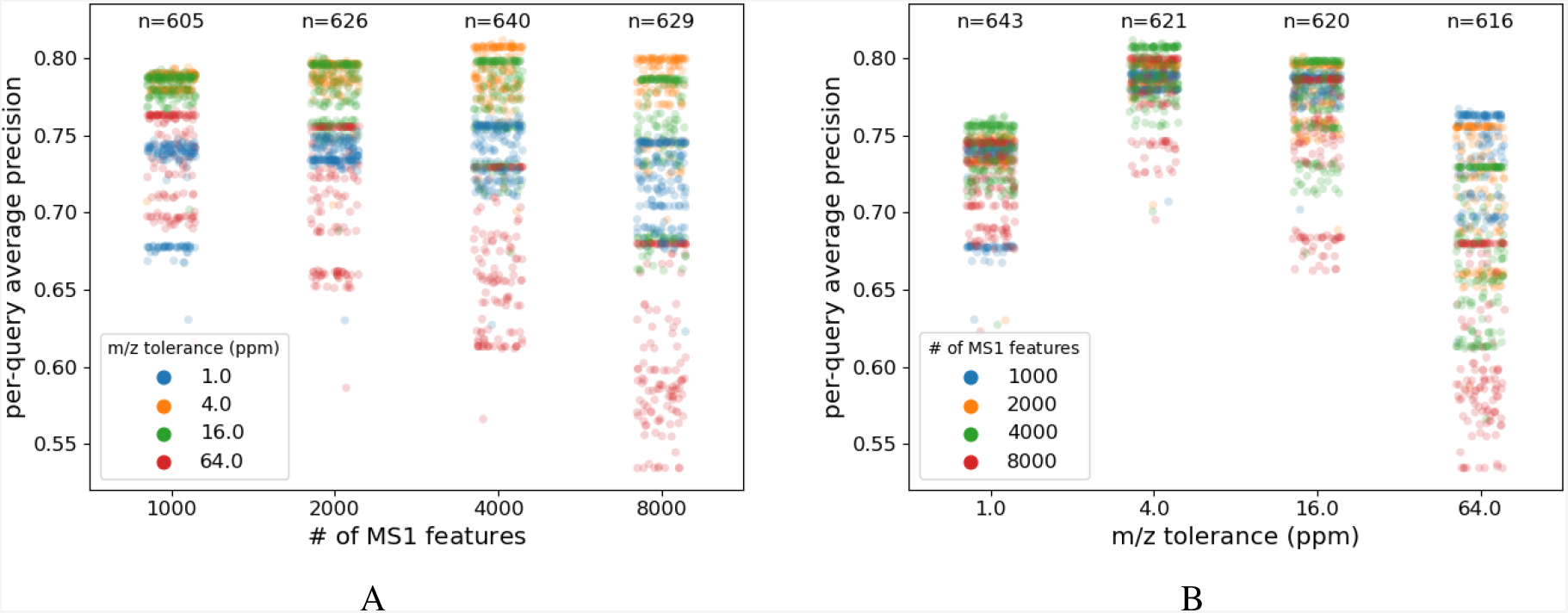
Performance as a function of by # of MS1 features or *m/z* tolerance. These figures plot the performance of 2500 different hyperparameterizations of MS1Connect as a function of (A) # or MS1 features or (B) *m/z* tolerance. The best performance occurs when *m/z* tolerance is set to 4 ppm and # of MS1 features is set to 4000. Note that the values near the top are the number of points found in each column.

Given that best performance occurred when *N* = 4000 and *m/z* tolerance = 4 ppm, we fixed these two hyperparameters and then asked whether there were any trends in the performance as a function of the remaining hyperparameters. We found that *λ*_4_ was correlated with performance (Supplemental Figure 8A). The performance increased as *λ*_4_ increased from 0.0 to 0.9, then performance decreased as *λ*_4_ approached 1.0. This trend suggested that larger *λ*_4_ may lead to higher performance and hence that supermodularity is important for achieving high performance. This is also shown by the fact that the best performing hyperparameterizations of MS1Connect occurred when *λ*_4_ = 0.9. In addition to *λ*_4_, *λ*_2_ was also found to be correlated with performance (Supplemental Figure 8B). In general, the performance MS1Connect decreased as *λ*_2_ increased from 0.0 to 1.0. This result indicated that *M*_2_, which measures intensity, is not useful for achieving high performance. However, we note that another hyperparameter, *N*, limits MS1 features in the bipartite graph to high-intensity features. None of the remaining hyperparameters were correlated with performance (Supplementary Figures 9–10).

## 5 Discussion

In this work, we introduced a new method, MS1Connect, that measures the similarity between pairs of proteomics runs. We showed evidence that MS1Connect successfully measures proteomics run similarity. Specifically, we showed that MS1Connect scores can be used for classifying what species a sample was generated from, and we showed evidence that MS1Connect scores can replicate database search results.

We also considered including in our empirical comparison a method based on dynamic time warping, but this approach turned out to be computationally infeasible. For this approach, we used FastDTW,^29^ with a cosine distance as the distance metric, to score the warping between two runs. We estimated that running dynamic time warping between all pairs of runs in our training dataset for the largest *m/z* bin width (0.01 Da) in our hyperparameter range would take approximately three weeks and 300 Gb of memory. The large memory requirement is due to the large matrices being compared to each other. For example, the run with the largest matrix had approximately 30,000 time bins and 700,000 *m/z* bins (*m/z* bin width of 0.0035 Da). The long time requirement resulted from having to calculate the warping between all pairs of runs (*>*13,000 warpings).

Since MS1Connect only uses MS1 data, it is agnostic to acquisition style. Therefore, MS1Connect can calculate the similarity between runs that were collected by DIA, DDA, or by targeted methods. While we did not specifically test this idea, we note that one set of five *S. aureus* runs in the training data was collected by DIA while the remaining runs were collected by DDA. Our results suggested that MS1Connect scores can measure the similarity between DIA and DDA runs. In turn, this ability to measure similarity is the first step for allowing future joint analysis of DDA and DIA data.

While we have shown that MS1Connect scores can be used to predict metadata labels of runs, future works needs to be conducted to determine the limits of our method. For example, none of the data used in this study was isobarically labeled. MS1Connect could be extended to include these types of samples. In addition, MS1Connect has not been tested on a wide variety of samples, such as fractionated samples. Additional work could be pursued to test the applicability of MS1Connect in these scenarios.

Finally, we note that while we developed MS1Connect for use in proteomics our method can be used to analyze data from other mass spectrometry-based fields such as metabolomics and lipidomics. Our method may be especially useful in these fields because MS2-based identifications are frequently missing. Therefore, MS2-based methods may not be as valuable. In the future, MS1Connect could be applied to data generated by metabolomics and lipidomics.

## 6 Acknowledgments

Some of the work was funded by NIH award R01GM121818. In addition, some of the research described in this paper was conducted under the Laboratory Directed Research and Development Program at Pacific Northwest National Laboratory, a multiprogram national laboratory operated by Battelle for the U.S. Department of Energy. Andy Lin is grateful for the support of the Linus Pauling Distinguished Postdoctoral Fellowship program. Pacific Northwest National Laboratory is a multiprogram national laboratory operated by Battelle Memorial Institute for the United States Department of Energy under contract DE-AC06-76RLO.

