## Supplementary figures and tables for "MS1Connect: a mass spectrometry run similarity measure"

### 1 Approach

#### 1.1 Representation of a mass spectrometry run

There are four values that are returned by pyOpenMS: $m/z$, intensity, charge, and retention time (in seconds). Prior to analysis, we replaced two of the three values with normalized versions thereof. For the retention time normalization, we first selected the top $N$ most intense MS1 features to represent an entire run, where $N$ is a hyperparameter of MS1Connect. Then, we normalized the retention times of the N features on a scale of zero to one based on the proportion of total ion current detected up to a given time. In addition, we removed MS1 features with a normalized retention time less than $0.05$ and greater than $0.95$. Finally, we scaled the intensities to range from zero to one by dividing by the maximum intensity among all MS1 features in the given run.

#### 1.2 Selecting the best matching

As previously noted, computing a maximization over an exponential number of edge subsets is generally intractable. However, our particular setup allows for an efficient computational of an approximate solution. First, we note that $M(A) = \sum_{e_1,e_2 \in A} m(e_1, e_2)$ is itself a function over subsets of edges, and that the function sums over the submatrix associated with edge set $A$. Given two subsets $A, B$, since it is the case that $m(e_1, e_2) \geq 0$ for all $e_1, e_2$, this function is said to be *supermodular* because it has the property of that $M(A) + M(B) \leq M(A \cup B) + M(A \cap B)$.

Second, we see that $\mathcal{E}_U$ and $\mathcal{E}_V$ correspond to a set of subsets of edges that have certain critical properties:

1. $\emptyset \in \mathcal{E}_U$,
2. if $B \in \mathcal{E}_U$ and $A \subseteq B$ then $A \in \mathcal{E}_U$, and
3. if $A, B \in \mathcal{E}_U$ with $|A| < |B|$, then $\exists b \in B \setminus A$ such that $A + b \in \mathcal{E}_U$.

These three properties define a matroid.[1,2] In fact, $\mathcal{E}_U$ and $\mathcal{E}_V$ are a particular kind of matroid, called a *partition matroid*. A partition matroid says that we partition the edges $E$ into disjoint blocks. For $A$ to be independent in the matroid means that $A$ must not intersect each block by more than a certain limit. In the present case, the blocks of edges are defined by the edges incident to each of the nodes $u \in U$ and $v \in V$. Thus, the constraint $A \in (\mathcal{E}_U \cap \mathcal{E}_V)$

*Corresponding author: Box 355065, Foege Building, S220B, 3720 15th Ave NE, Seattle, WA 98195-5065, means that for $A$ to be feasible, it must simultaneously be a member of the independent sets of two partition matroids $\mathcal{E}_U$ and $\mathcal{E}_V$. Hence, to produce our run-pair score, we must solve an instance of supermodular maximization subject to two matroid constraints.

Ordinarily, computing the solution to such a problem is inapproximable, because supermodular maximization subject to even simple constraints in general is very hard and even hard to approximate.[3,4] However, recent work has shown that if the supermodular function exhibits certain properties, then an approximation algorithm is possible using a greedy procedure.[5] Ordinarily, the greedy algorithm can do arbitrarily poorly when the function is supermodular (see[4] for a simple example). However, when the function has limited curvature, an approximation is possible. We define the supermodular curvature as $\kappa^f = 1 - \min_{e \in E} f(e)/f(e \setminus E)$. It has been showed that $0 \leq \kappa^f \leq 1$ and if $\kappa^f = 1$, then constrained supermodular maximization is indeed inapproximable.[5] However, if $\kappa^f < 1$, then the greedy algorithm achieves an approximation bound of $1 - \kappa^f$, which means that $f(\tilde{S}) \geq (1 - \kappa^f) f(S^*)$ where $\tilde{S}$ is the solution provided by the greedy algorithm, and $S^*$ is the optimal solution.

The greedy algorithm is fairly simple. We start with $S \leftarrow \emptyset$ and we then repeat the step

$$S \leftarrow S \cup \operatorname{argmax}_{e \in E \setminus S} f(S + e)$$

until the constraints are no longer satisfied. After this procedure finishes, we multiply in the de-normalization term $(\sum_{i=1}^{4} \lambda_i M_i(E))$ as a post-processing step. This algorithm runs quickly and scalably even with very large sets $|E|$, and we therefore use this procedure for computing the score in Equation 1.

The curvature of $f$ is very dependent on the matrix $M$. It turns out that as long as the diagonal entries of $M$ are non-zero, then $\kappa^g < 1$. In Supplementary Figure 4, we show that our best results correspond to the case where the diagonal is non-zero, and this happens as long as $\lambda_4 < 1$ in Equation (4), and thus $\kappa^g < 1$. These observations offer mathematical justification for the performance of our algorithm.

### 2 Methods

#### 2.1 Hyperparameter search

MS1Connect ($M_1$ – $M_4$) has nine different hyperparameters (Table 1). Two of them affect the generation of the bipartite graph while the remaining seven hyperparameters affect the scoring of a specific selected edge set. The first hyperparameter, $N$, is the maximum number of MS1 features used, in each run, to generate the bipartite graph. The next hyperparameter $\delta_1$ is the $m/z$ threshold (in ppm) used for connecting two MS1 features. Each of the four $\lambda$ hyperparameters are used to weight each of the four terms in the objective function. Finally, $\alpha$, $\beta$, and $\gamma$ affect the rate at which the exponential functions decrease in $M_3$ and $M_4$.

To determine the best performing hyperparameter set, as measured by QAP, we sampled the hyperparameter space using a random grid search. In a random grid search, the range and values of each hyperparameter is predetermined. Then, for each instance of a hyperparameter set, random values are selected from the grid. Our hyperparameter grid consists of 954,976 possible sets of hyperparameters (Supplemental Table 1). The number of MS1 features used in the bipartite graph generation ranged from 1,000 to 4,000 with two fold-change increments. The $m/z$ tolerance ranged from 1 ppm to 16 ppm with four-fold change increments. All four $\lambda$ hyperparameters were allowed to range between zero and one, inclusive, with linear increments of 0.1, subject to the constraint that $\sum_1^4 \lambda_i = 1.0$. Finally, the hyperparameters $\alpha$, $\beta$, $\gamma$ ranged from $10^{-6}$ to 1.0, with 10-fold change increments. We evaluated a total of 1,716 different randomly chosen hyperparameter sets.

In addition to a hyperparameter search for MS1Connect ($M_1$ – $M_4$), we conducted an exhaustive hyperparameter search for each of the four versions of MS1Connect that only used a single term. Each version used the same hyperparameter grid as MS1Connect ($M_1$ – $M_4$) except that we added two additional values (250 and 500) to the number of MS1 features for the variants of MS1Connect that only use $M_1$, $M_2$, or $M_3$. We note that not all versions of MS1Connect use all the previously described hyperparameters. For example, the only relevant hyperparameters in MS1Connect ($M_1$ only) are number of MS1 features and $m/z$ tolerance. The best performing set of hyperparameters for each method can be found in Supplemental Table 4. Note that MS1Connect ($M_4$ only) has six different sets of hyperparameters that yield the best performance.

Finally, we performed an exhaustive hyperparameter search for our eight baselines (Supplemental Table 3). These baselines have two different hyperparameters: number of MS1 features and the $m/z$ bin width $\delta_2$. The hyperparameter grid for these methods allowed $\delta_2$ to range from 0.0025 to 0.01 Da with $\sqrt{2}$-fold change increments, and the number

| hyperparameter | possible values |
|---|---|
| # of MS1 features | 1000, 2000, 4000, 8000 |
| $m/z$ tolerance (ppm) | 1, 4, 16, 64 |
| $\alpha$ | $10^{-6}$, $10^{-5}$,..., 1.0 |
| $\beta$ | $10^{-6}$, $10^{-5}$,..., 1.0 |
| $\gamma$ | $10^{-6}$, $10^{-5}$,..., 1.0 |
| $\lambda_1$ | 0.0, 0.1,..., 1.0 |
| $\lambda_2$ | 0.0, 0.1,..., 1.0 |
| $\lambda_3$ | 0.0, 0.1,..., 1.0 |
| $\lambda_4$ | 0.0, 0.1,..., 1.0 |

Supplemental Table 1: **MS1Connect hyperparameter search grid.** Table of the grid that was searched during the hyperparameter search. We conducted an exhaustive hyperparameter search for each version of MS1Connect that only considered a single term, which yielded 784 hyperparameter sets. In addition, we evaluated 1,716 randomly chosen sets of hyperparameters. Altogether, we evaluated 2,500 out of the 954,976 possible hyperparameter sets.

of MS1 features ranged from 250 to 8,000 with two-fold change increments. The best set of hyperparameters for each method can be found in Supplemental Table 5.

#### 2.2 Microbial cultivation

Microbial strains used in this study were: *Escherichia coli* 15597, *Salmonella enterica* serovar Typhimurium 14028, *Bacillus cereus* 14579, *Bacillus thuringiensis* Al Hakam, *Bacillus thuringiensis* HD600 and *Porphyrobacter* sp. HL-46. *Escherichia coli* 15597, *S. enterica* serovar Typhimurium 14028, and *B. cereus* 14579 were cultivated on Tryptic Soy Agar (TSA) from glycerol stocks and then into Tryptic Soy Broth for 16 hours at 37 °C. *Bacillus thuringiensis* Al Hakam, and *B. thuringiensis* HD600 were grown at 32 °C in TSB, and *Porphyrobacter* sp. HL-46 was grown in Hot Lake autotroph media as described in Cole et al.[6] Cultures were centrifuged to pellet cellular biomass and washed once in sterile phosphate buffered saline (PBS). PBS was removed and the pellets were stored at -80 °C until prepared for peptide analysis.

#### 2.3 Microbial sample preparation

Bacterial cell pellets were thawed and resuspended in a lysis buffer containing 6 M urea and 14.3 mM betamercaptoethanol in 50 mM Tris-HCl pH 8. Samples were incubated at 60 °C in a thermomixer with gentle shaking (300 rpm) for one hour, then 50 mM ammonium bicarbonate was added to reduce the urea concentration 10 fold. Trypsin Gold (Promega) was resuspended in acetic acid to 1 $\mu$g/$\mu$L concentration and 2 $\mu$L was added to each sample. Samples were incubated overnight at 37 °C in a thermomixer with gentle shaking (300 rpm). Digested peptide samples were spun down briefly to pellet any remaining solid debris and the supernatant was subjected to solid phase extraction (SPE). SPE cartridges (Phenomenex; Strata C18-T (55 $\mu$m, 140 Å), 100 mg/1 mL, cat # 8B-S004-EAK) were loaded into the vacuum manifold and samples were processed according to the manufacturers recommendation. Briefly, cartridges were conditioned with 1 mL methanol and washed with 1 mL 0.1% trifluoroacetic acid (TFA) in water. The sample was added to the cartridge, followed by a wash with 1 mL 5% acetonitrile/95% 0.1% TFA in water. Finally, the sample was eluted with 1 mL 80% acetonitrile and 20% 0.1% TFA in water. Samples were concentrated to near dryness down using a SpeedVac and resuspended in 30 $\mu$L 0.1% TFA water. A BCA assay (Pierce) was performed to determine peptide concentration. Samples were diluted to 0.1 $\mu$g/$\mu$L and stored at -20 °C until MS analysis.

#### 2.4 Plant sample processing and preparation

In addition to the bacterial samples used in this study, plant material from *Nicotiana benthamiana* was also used. Leaf plant material was processed and prepared as previously described.[7] Briefly, material was washed, frozen in liquid nitrogen, and ground in a precooled mortar in the presence of liquid nitrogen. Ground plant material (150 mg) was precipitated at -20°C overnight in 10% trichloroacetic acid (TCA) with 0.07% betamercaptoethanol in cold acetone. The next day, the sample was centrifuged at 10,000 × g for 15 minutes, the supernatant was removed, and the remaining pellet was washed twice with ice cold acetone with 0.07% betamercaptoethanol. The pellet was then resuspended in the lysis buffer as described for microbial cell samples and all downstream reduction, denaturation, and digestion steps were the same for the plant sample as for the microbial cell samples.

#### 2.5 Liquid chromatography-tandem mass spectrometry

**Lab 1.** Liquid chromatography separation was performed using an Agilent 1200 HPLC instrument with a 40 cm long 0.15 mm ID fused silica packed with Jupiter 5 $\mu$m C-18 resin. Mobile phase A was prepared with 5% acetonitrile and 0.1% formic acid in nano-pure H2O; mobile phase B was prepared with 95% acetonitrile and 0.1% formic acid in nano-pure H2O. The flow rate was 2 $\mu$L per minute with a reversed phase gradient transitioning from 0% solution B to 45% solution B over the course of 60 min for separation followed by a wash and regeneration step. An Orbitrap XL mass spectrometer (Thermo Electron, Thousand Oaks, CA) was used to analyze the eluate from the HPLC, which was directly ionized and transferred into the gas phase with an electrospray emitter (operated at 3.5 kV relative to the mass spectrometer interface). The ion transfer tube on the Orbitrap system was maintained at 200 °C and 200 V with an ion injection time set for automatic gain control with a maximum injection time of 200 ms for $5 \times 10^7$ charges in the linear ion trap. Ion selection was achieved using dynamic parent ion selection in which the five most abundant ions were selected for MS/MS using a 3 m/z window. Each sample was analyzed in technical triplicate.
**Lab 2.** A Waters nano-Acquity dual pumping UPLC system (Milford, MA) was configured for on-line trapping of a 5 $\mu$L injection at 5 $\mu$L/min with reverse-flow elution onto the analytical column at 300 nL/min. Columns were packed in-house using 360 µm o.d. fused silica (Polymicro Technologies Inc., Phoenix, AZ) with 2-mm sol-gel frits for media retention and contained Jupiter C18 media (Phenomenex, Torrence, CA) in 5$\mu$m particle size for the trapping column (150$\mu$m i.d. x 4cm long), with 3$\mu$m particle size for the analytical column (75 $\mu$m i.d. x 70 cm long). Mobile phases consisted of (A) 0.1% formic acid in water and (B) 0.1% formic acid in acetonitrile with the following gradient profile (min, %B): 0,1; 8,1; 10,8; 28,12; 83,30; 105,45; 108,95; 118,95; 122,50; 124,95; 126,1; 128,50; 130,50; 132,1; 152,1.

MS analysis was performed using an Q Exactive HF mass spectrometer (Thermo Scientific, San Jose, CA) outfitted with a home-made nano-electrospray ionization interface. Electrospray emitters were prepared using 150 $\mu$m o.d. x 20 um i.d. chemically etched fused silica.[8] The ion transfer tube temperature and spray voltage were 325ºC and 2.3 kV, respectively. Data were collected for 100 min following a 20 min delay from sample injection. FT-MS spectra were acquired from 400-2000 m/z at a resolution of 60k (AGC target 3e6) and while the top 12 FT-HCD-MS/MS spectra were acquired in data dependent mode with an isolation window of 2.0 m/z and at a resolution of 15k (AGC target 1e5) using a normalized collision energy of 30 and a 45 sec exclusion time.

#### 2.6 Database search

A database search was conducted using Crux version 3.2[9,10] on the runs from the multi-species dataset using the combined p-value score function.[11] The spectra were searched against a concatenated database containing the proteomes of the following species: *Porphyrobacter* sp. YT40 (`https://www.uniprot.org/proteomes/UP000315943`), *Salmonella enterica* (`https://www.uniprot.org/proteomes/UP000054420`), *Escherichia coli* (`https://www.uniprot.org/proteomes/UP000000558`), *Bacillus cereus* (`https://www.uniprot.org/proteomes/UP000001417`), *Bacillus thuringiensis* (`https://www.uniprot.org/proteomes/UP000032057`), and *Nicotiana tabacum* (`https://www.uniprot.org/proteomes/UP000084051`). The proteomes were downloaded from Uniprot[12] in April 2021. The protein database was digested into peptides using the tide-index tool, and the search was performed by the tide-search tool. All parameters were set to their default values except that "mz-bin-width" was set to 1.0005079, "score-function" was set to "both", "exact-p-value" was set to "True", and "top-match" was set to one.

The spectra from the *Bacillus* genus data were searched in the same manner as the multi-species dataset except that the spectra were searched against a concatenated database containing the following species: *B. cereus* (`https://www.uniprot.org/proteomes/UP000001417`), *B. cytotoxicus* (`https://www.uniprot.org/proteomes/UP000002300`), *B. mycoides* (`https://www.uniprot.org/proteomes/UP000001754`), *B. pseudomycoides* (`https://www.uniprot.org/proteomes/UP000001378`), *B. thuringiensis* (`https://www.uniprot.org/proteomes/UP000011719`), *B. weihenstephanensis* (`https://www.uniprot.org/proteomes/UP000002154`), and *B. toyonensis* (`https://www.uniprot.org/proteomes/UP000017860`). These proteomes were downloaded from Uniprot[12] in January of 2022.

#### 2.7 Data

**Species training dataset.** A total of 166 RAW files from 33 PRIDE IDs were downloaded from PRIDE in November of 2020. These 166 runs each originated from a sample that contains one of nine different species (Supplemental

Figure 2A). For this dataset we used data from up to five PRIDE IDs per species, except that human had six PRIDE IDs, and downloaded up to five RAW files per PRIDE ID. The mass spectrometry runs were collected using various types of Thermo Scientific instrumentation (San Jose, CA). We did not include any off-line fractionated samples or labeled samples (*e.g.* TMT or ITRAQ). All samples were digested using trypsin. Additional information about this dataset can be found in Supplemental File 1.

**Species test dataset.** An additional 130 RAW files from 23 PRIDE IDs were downloaded from PRIDE in December of 2020 for compilation into a species test dataset. Each run consists of data from one of nine different species (Supplemental Figure 2B). Like the training dataset, we used five RAW files per PRIDE ID; however, we only used data from three different PRIDE IDs per species. One exception is that we only collected *Plasmodium falciparum* data from two different PRIDE IDs. The other exception is that all 15 runs of the *Yersinia pestis* data came from a single PRIDE ID. However, the data from that project itself is an aggregation of several different experiments. None of the PRIDE IDs nor runs in this dataset overlap with the data in the species training dataset. Additional information about this dataset can be found in Supplemental File 2.

**Multi-species data.** This dataset consists of 16 different samples (labeled A-P) that were each injected into a mass spectrometer four times, which resulted in 63 runs (sample P was run three times). Each sample contains up to three different organisms from a set of seven possible species (Supplemental Table 6). If a sample contained multiple species then they were mixed together in equal concentrations. Each sample was injected into an LTQ-Orbitrap XL three times and one time into an Q-Exactive. We note that the LTQ-Orbitrap XL and the Q-Exactive instruments are from different laboratories and therefore have different liquid chromatographies and instrument parameters. Additional information about this dataset can be found Supplemental File 3, and the raw files have been deposited into PRIDE with dataset identifier PXD027791.

***Bacillus* genus data.** A total of 42 RAW files were downloaded from PRIDE (PXD003669) in November of 2021.[13] These runs originated from seven different strains of the *Bacillus cereus* group. The seven strains analyzed were *B. cereus* DSM 31, *B. cytotoxicus* DSM 22905, *B. mycoides* DSM 2048, *B. pseudomycoides* DSM 12442, *B. thuringiensis* DSM 2046, *B. weihenstephanensis* DSM 11821, and *B. toyonensis* CECT 876. A total of six runs, two technical replicates of three independent biological replicates, for each strain was run on a nanoAcquity UPLC (Waters Inc., Milford, USA) connected to a Q-Exactive mass spectrometer (Thermo Scientific). Additional information about this dataset can be found Supplemental File 4.

### 3 Note

#### 3.1 Explanation why supermodular methods outperform modular methods

Our work showed evidence that supermodular methods outperform modular methods. Here we speculate why modular terms may be insufficient for measuring similarities between mass spectrometry runs by discussing the modular $M_3$ term. A single edge will score highly with $M_3$ when the normalized retention time between the two features that make up an edge are very similar. At first glance, this seems reasonable because two peptide features with the same $m/z$ and retention time should be the same ion. However, this observation fails to account for systematic differences in retention time that result from differing liquid chromatography conditions. Therefore, while we expect the order in which peptides elute off the column to approximately match between two runs, we do not expect such correspondence in retention time. As a result, an edge with a large retention time shift could be correct. Using supermodularity to consider pairs of edges overcomes this issue because it considers a neighborhood of edges. If a set of edges in the same region have the same retention time shift, it indicates that a systemic time shift has occurred and the retention time shifts are correct.

| Variable | Definition |
|---|---|
| $G = (U, V, E)$ | Bipartite graph containing two vertex sets $U$ and $V$ and edge set $E$ |
| $U$ | vertex set of MS1 features from one run |
| $V$ | vertex set of MS1 features from a second run |
| $r$ | mass spectrometry run |
| $E$ | set of valid edges |
| $u$ | node in $U$ |
| $v$ | node in $V$ |
| $e$ | instance of valid edge |
| $s(e)$ | normalized retention time shift of edge e |
| $u(e)$ | MS1 feature $u$ that is a part of edge $e$ |
| $v(e)$ | MS1 feature $v$ that is a part of edge $e$ |
| $t(u)$ | normalized retention time of MS1 feature $u$ |
| $m(u)$ | $m/z$ of MS1 feature $u$ |
| $I(u)$ | normalized intensity of $u$ |
| $c(u)$ | charge of MS1 feature $u$ |
| $A$ | set of edges scored by objective function |
| $S$ | objective function used to a score a set of edges |
| $\mathcal{E}_V$ | degree constraint on nodes in $V$ |
| $\mathcal{E}_U$ | degree constraint on nodes in $U$ |
| $\delta_1$ | $m/z$ tolerance in ppm |
| $\delta_2$ | $m/z$ bin width in Daltons |
| $N$ | the number of most intense MS1 features used to represent a run |
| $n$ | number of retention time bins |
| $M_i$ | one of four terms in the objective function |
| $\lambda_i$ | hyperparameter that weights the importance of each of the four terms in the objective function |
| $\alpha$ | hyperparameter that is used in $M_3$ |
| $\beta$ | hyperparameter that is used in $M_4$ |
| $\gamma$ | hyperparameter that is used in $M_4$ |
| $\kappa^f$ | supermodular curvature |
| $P_k$ | precision at $k$ |
| $R_k$ | recall at $k$ |
| $q$ | query run |
| $R$ | set of repository runs |
| $C$ | number of repository runs |
| $X$ | matrix representation of mass spectrometry run |
| $Y$ | matrix representation of mass spectrometry run |
| $G$ | function that is either the product, maximum, minimum, or indicator of two input values |

Supplemental Table 2: **Notation.** Notation used in this manuscript.

| hyperparameter | hyperparameter range |
|---|---|
| # of MS1 features | 250, 500, 1000, 2000, 4000, 8000 |
| $m/z$ bin width ($\delta_2$) | 0.0025, 0.0035, 0.005, 0.0071, 0.01 |
| retention time bins | 1, 2, 4, 8 |

Supplemental Table 3: **Hyperparameter search grid for baselines.** Table of the grid that was searched during the exhaustive hyperparameter search for each of the baselines.

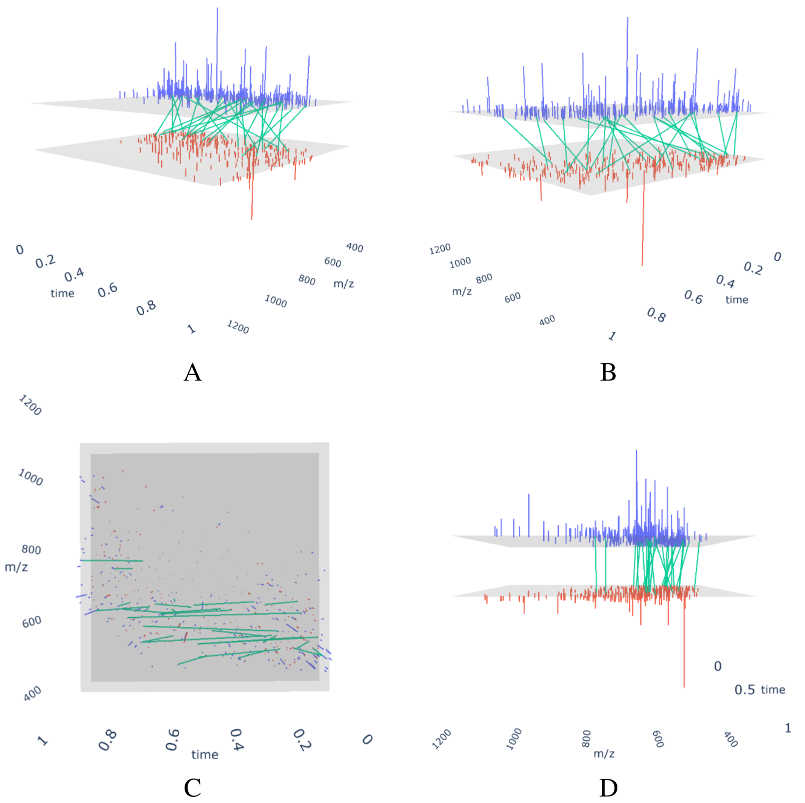


Supplemental Figure 1: **Views of bipartite graph.** Four views of the bipartite graph formed by a pair of mass spectrometry runs. The purples lines are the MS1 features from one run (006_EC-D2O_A17.raw) and the orange lines are the MS1 features from the second run (11B_RINICHI_sTgw10.raw). The green lines represent the set of all possible edges $E$ that link MS1 features with similar $m/z$. The three axes shown in these plots are $m/z$ retention time, and intensity. The two gray planes represent the bottom of each run, with respect to intensity. Note in D how edges are nearly vertical and only link MS1 features with similar $m/z$.

| Method | # MS1 features | $m/z$ tolerance (ppm) | $\lambda_1$ | $\lambda_2$ | $\lambda_3$ | $\lambda_4$ | $\alpha$ | $\beta$ | $\gamma$ |
|---|---|---|---|---|---|---|---|---|---|
| MS1Connect ($M_1$ – $M_4$) | 4000 | 4 | 0.0 | 0.1 | 0.0 | 0.9 | 0.1 | $10^{-5}$ | 1.0 |
| MS1Connect ($M_1$ only) | 1000 | 4 | 1.0 | 0.0 | 0.0 | 0.0 | NA | NA | NA |
| MS1Connect ($M_2$ only) | 500 | 16 | 0.0 | 1.0 | 0.0 | 0.0 | NA | NA | NA |
| MS1Connect ($M_3$ only) | 1000 | 4 | 0.0 | 0.0 | 1.0 | 0.0 | $10^{-5}$ | NA | NA |
| MS1Connect ($M_4$ only)* | 4000 | 4 | 0.0 | 0.0 | 0.0 | 1.0 | NA | 1.0 | 0.001 |

Supplemental Table 4: **Best performing MS1Connect hyperparameters.** The best performing set of hyperparameters for each MS1Connect method. *Six different hyperparameterizations of MS1Connect (M4) have the best performance. Specifically, the best performance occurs when $\beta$ and $\gamma$ are 0.1 and $10^{-5}$, 0.1 and $10^{-6}$, 1.0 and $10^{-3}$, 1.0 and $10^{-4}$, 1.0 and $10^{-5}$, or 1.0 and $10^{-6}$ respectively.

| Method | type | # MS1 features | $m/z$ tolerance (Da) | # RT bins (n) |
|---|---|---|---|---|
| # $m/z$ bins in common | indicator | 1000 | 0.0071 | NA |
| # $m/z$ + RT bins in common | indicator | 1000 | 0.0050 | 2 |
| # $m/z$ bins in common | count | 500 | 0.0071 | NA |
| # $m/z$ + RT bins in common | count | 1000 | 0.0050 | 2 |
| # $m/z$ bins in common | min | 1000 | 0.0071 | NA |
| # $m/z$ + RT bins in common | min | 1000 | 0.005 | 2 |

Supplemental Table 5: **Best performing baseline hyperparameters.** The best performing set of hyperparameters for each baseline method. We did not include the best set of hyperparameters for the max version of the baseline as it had poor performance.

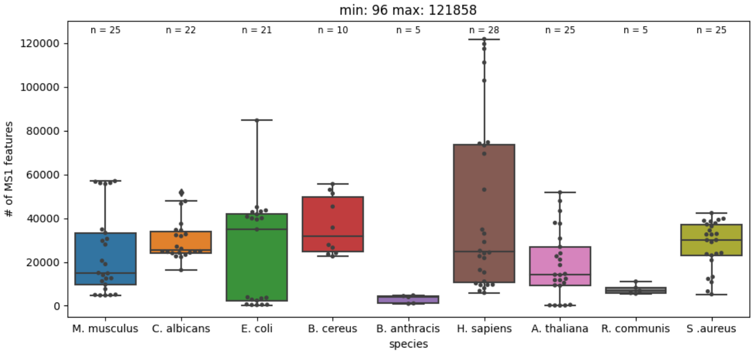


A

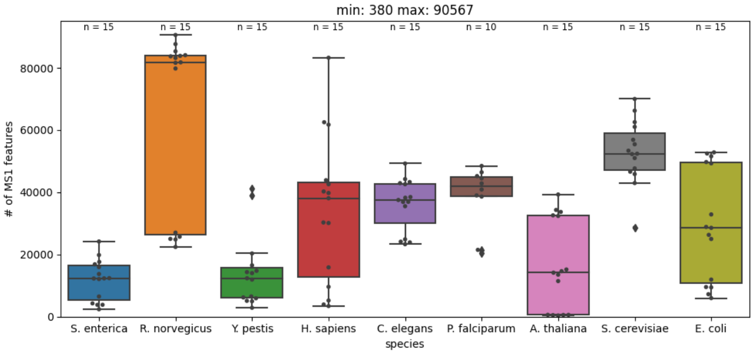


B

Supplemental Figure 2: **Number of MS1 features per run.** A series of boxplots of the number of MS1 features per run split by species for the (A) species training data and (B) the species test data. In addition to the boxplots, each point shows the number of MS1 features per run. The text near the top of the plot indicates the number of runs that each label has.

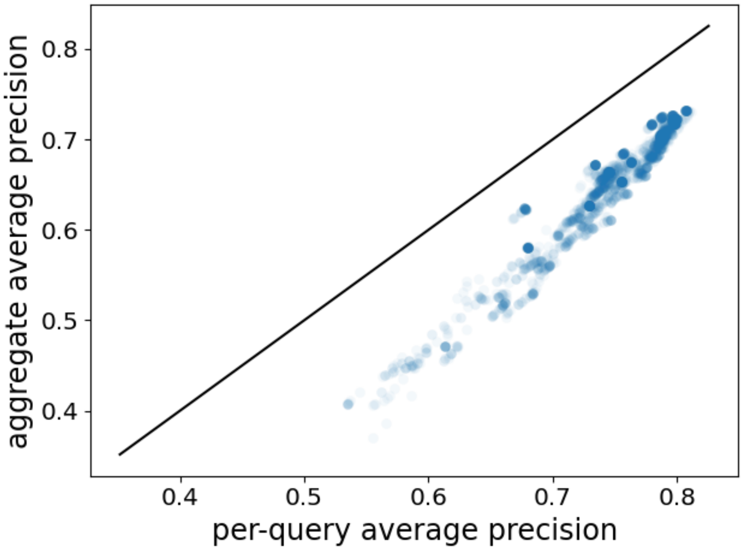


Supplemental Figure 3: **Comparison of per-query average precision and aggregate average precisio.** Scatter plot of the per-query average precision and aggregate average precision of the species training dataset over 2,500 different hyperparameterizations. These metrics are highly correlated with a Pearson correlation of 0.9689. The solid black line shows the $y = x$ line.

| Sample ID | Porphyrobacter | *S. enterica* | *E. coli* | *B. cereus* | *B. thuringiensis* Al Hakam | *B. thuringiensis* HD600 | *N. benthamiana* |
|---|---|---|---|---|---|---|---|
| A | ✓ | | | | | | |
| B | | ✓ | | ✓ | | | |
| C | | | ✓ | | | | |
| D | | | | ✓ | | | |
| E | | | | | ✓ | | ✓ |
| F | | | | | | ✓ | |
| G | ✓ | | | | ✓ | ✓ | |
| H | | | ✓ | ✓ | | | ✓ |
| I | ✓ | | | | | | ✓ |
| J | ✓ | | ✓ | | | | |
| K | | ✓ | | | | | |
| L | | | | | ✓ | ✓ | |
| M | | | | | ✓ | | |
| N | | | | ✓ | | ✓ | |
| O | | ✓ | ✓ | | | | |
| P | | | | | | | ✓ |

Supplemental Table 6: **Table of species data.** A table of the species found in each sample. Note that *B. cereus* is *Bacillus cereus* 14579, *S. enterica* is *Salmonella enterica* Typhimurium strain 14028, and *E. coli* is *Escherichia coli* 15597. In addition, *B. cereus*, *B. thuringiensis* Al Hakam, and *B. thuringiensis* HD600 are often considered the same species. Horizontal lines are used visualization purposes.

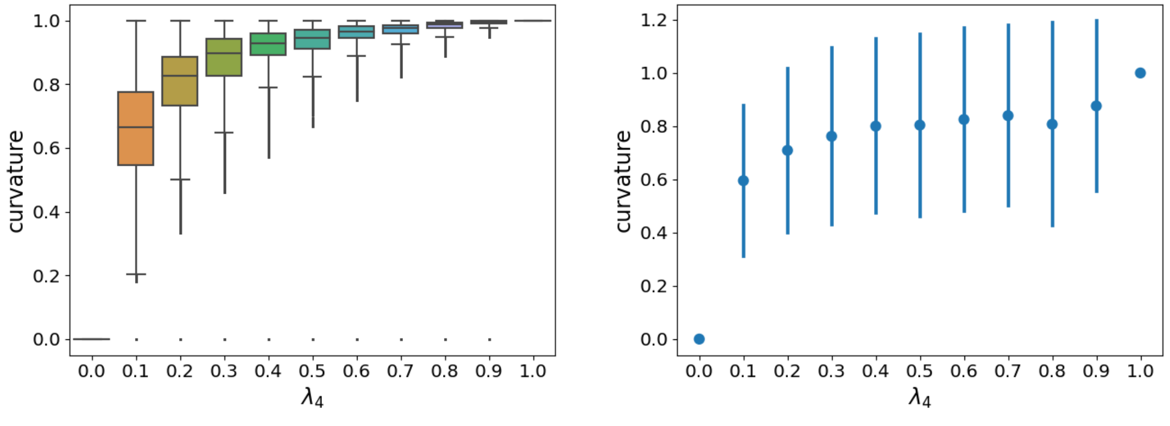


A B

Supplemental Figure 4: **Curvature as a function of $\lambda_4$.** These plots show the curvature value as a function of $\lambda_4$. The best performance of MS1Connect occurs when $\lambda_4 = 0.9$ A) A boxplot of the curvature values showing mean and inter-quartile range. B) A plot of the mean and standard deviation of the curvature values. Note that curvature values, by definition, range from zero to one, inclusive.

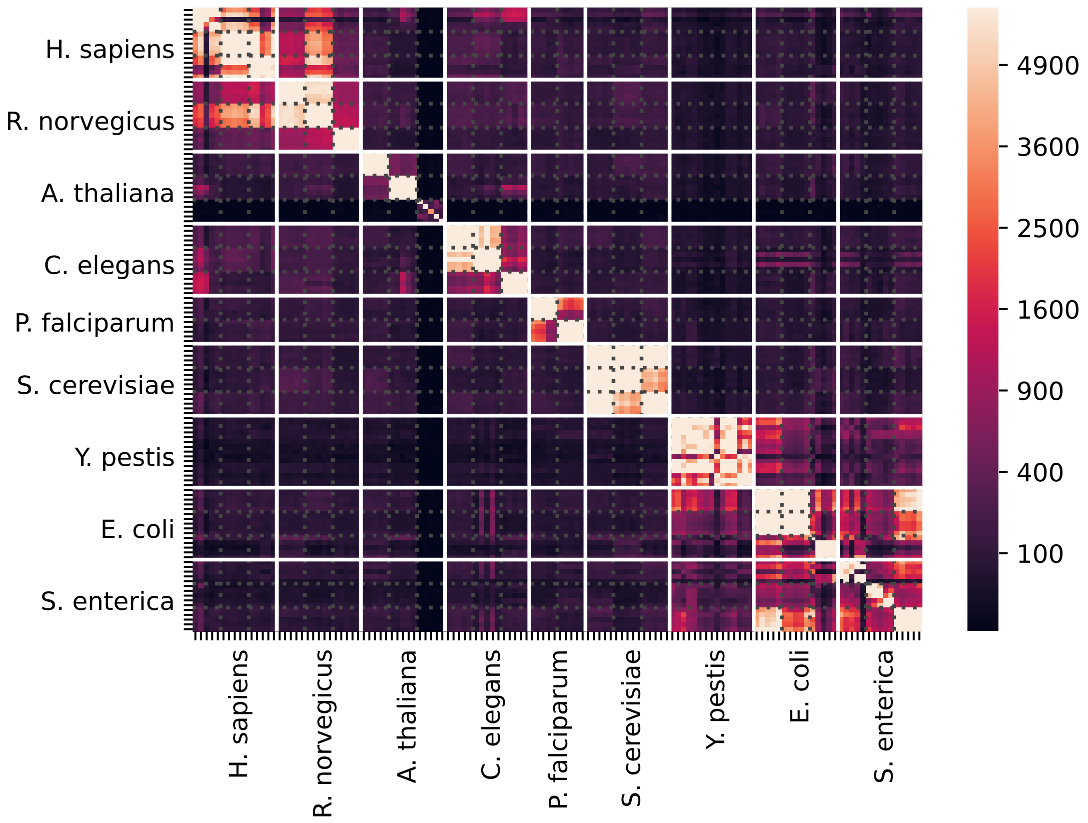


Supplemental Figure 5: **Heatmap of MS1Connect scores for species test data.** Each cell is colored by the MS1Connect score between a pair of runs. The solid white lines denote the border between different species while the dotted gray lines delineate the border between different experiments (PRIDE ID).

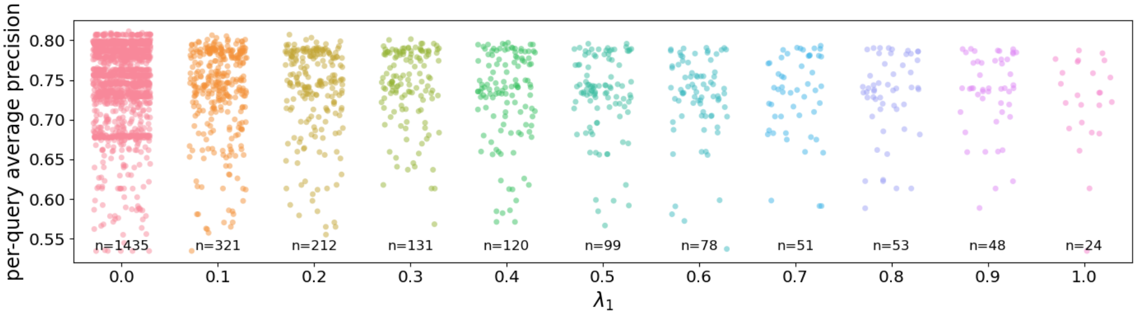


A

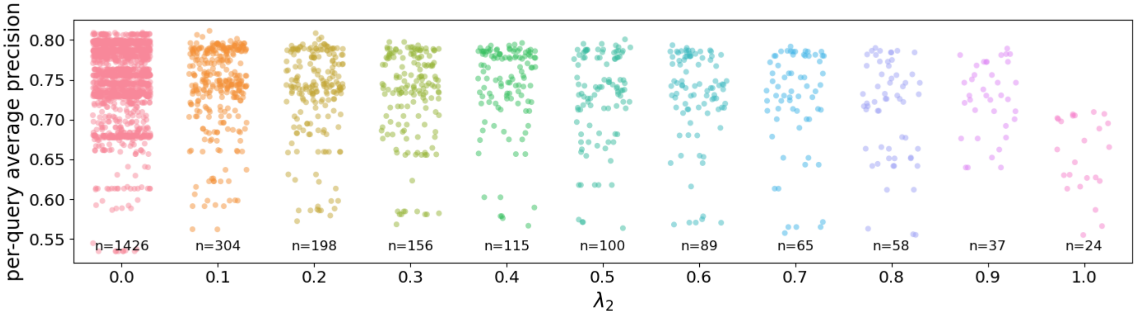


B

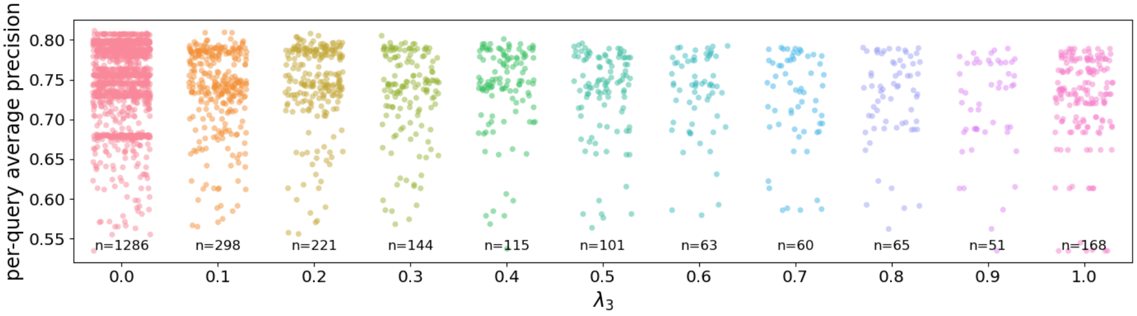


C

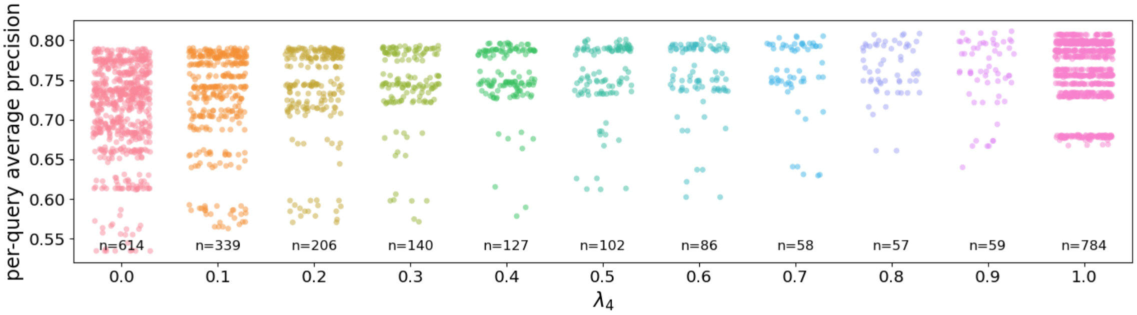


D

Supplemental Figure 6: **Performance as a function of $\lambda$.** This figures plots the performance of 2500 different hyperparameterizations of MS1Connect as a function of (A) $\lambda_1$, (B) $\lambda_2$, (C) $\lambda_3$, (D) or $\lambda_4$. Note that the values near the bottom are the number of points found in each column.

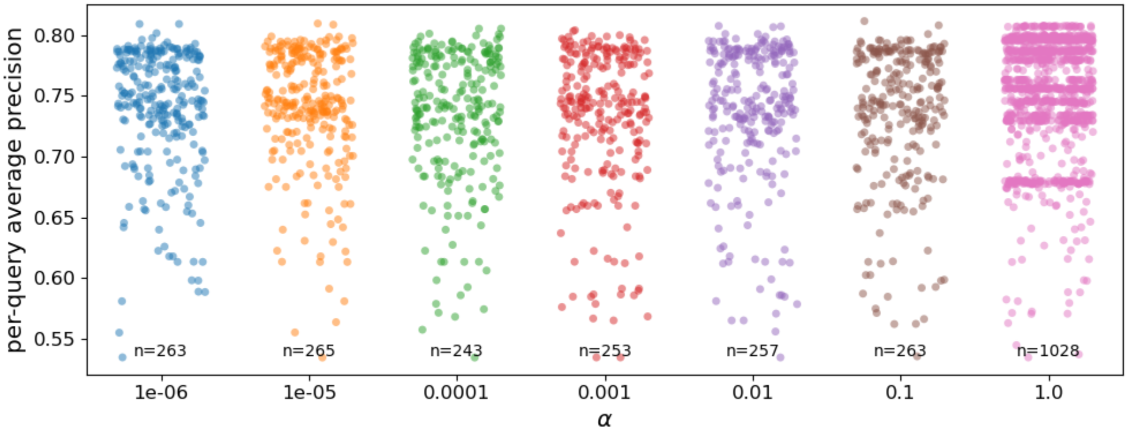


A

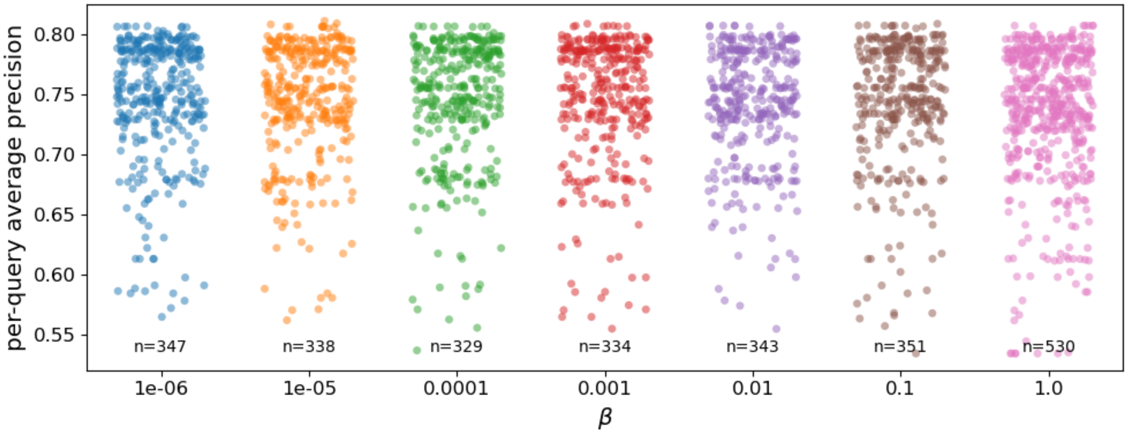


B

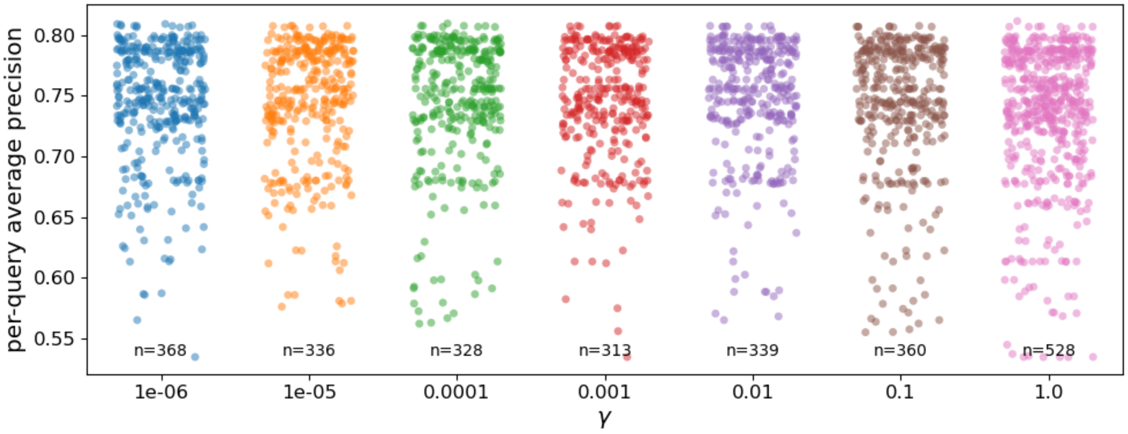


C

Supplemental Figure 7: **Performance as a function of $\alpha$, $\beta$, or $\gamma$.** This figures plots the performance of 2500 different hyperparameterizations of MS1Connect as a function of (A) $\alpha$, (B) $\beta$, (C) or $\gamma$. Note that the values near the bottom are the number of points found in each column.

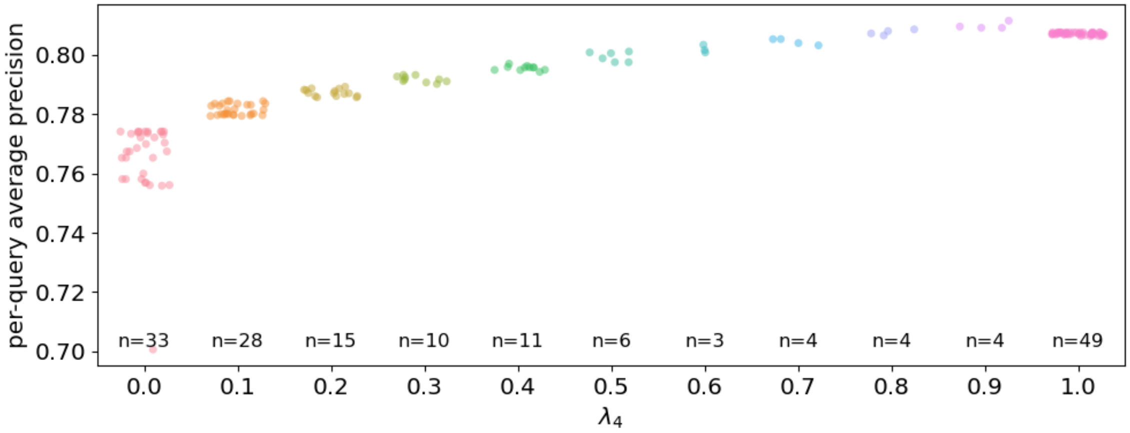


A

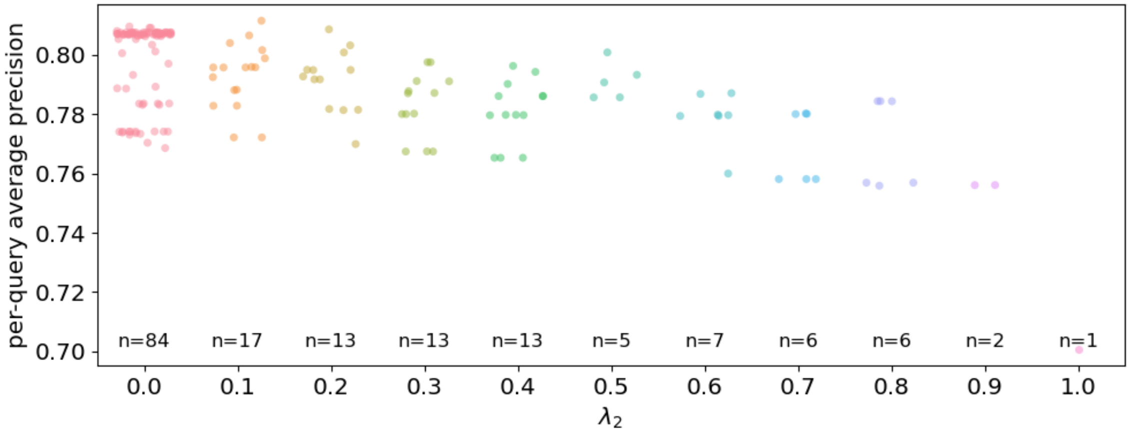


B

Supplemental Figure 8: **Performance of various MS1Connect hyperparameterizations after fixing $N$ and $\delta_1$.** Each panel plots plots the performance of MS1Connect as a function of (A) $\lambda_4$ or (B) $\lambda_2$. Performance tends to decrease as $\lambda_2$ increases from 0.0 to 1.0. In addition, performance tends to increase as $\lambda_4$ increases from 0.0 to 0.9 and then decreases as $\lambda_4$ increases from 0.9 to 1.0. Note that the values near the bottom are the number of points found in each column. For both plots, there tends to be fewer points as the $\lambda$ value increases because $\sum_1^4 \lambda_i = 1.0$. In addition, there are fewer points when $\lambda_2$ is large because $M_2$ does not use hyperparameters $\alpha$, $\beta$, and $\gamma$.

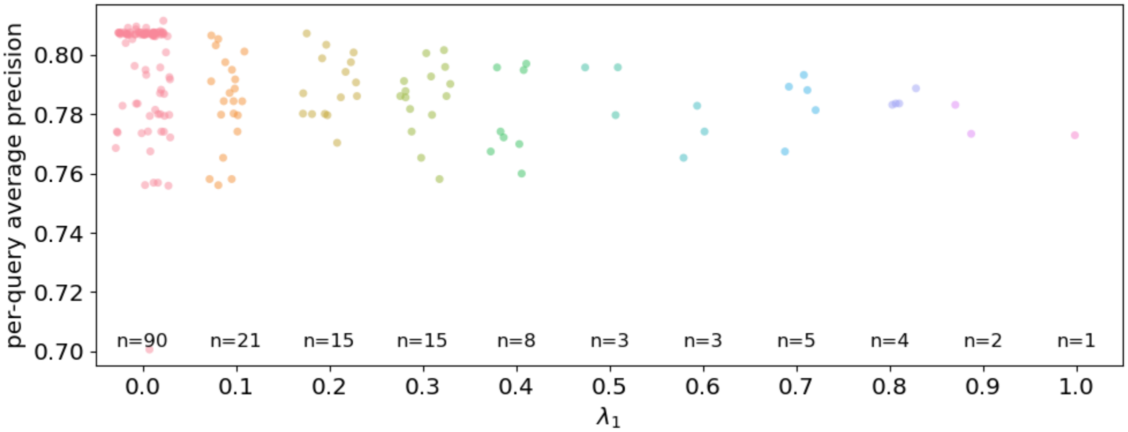


A

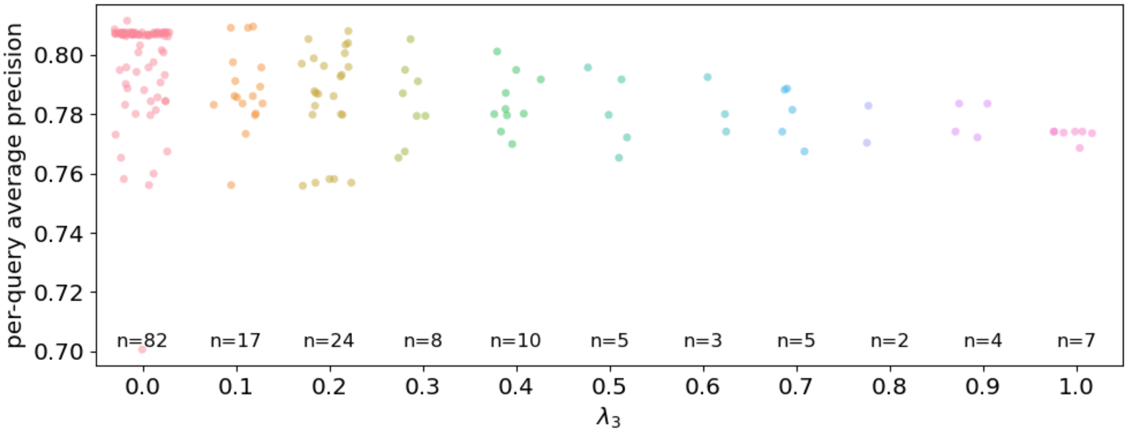


B

Supplemental Figure 9: **Performance as a function of $\lambda_1$ or $\lambda_3$.** This figures plots the performance of MS1Connect, given $N = 4000$ and $\delta_1 = 4$ ppm, as a function of (A) $\lambda_1$ (B) or $\lambda_3$. Note that the values near the bottom are the number of points found in each column. For both plots, there tends to be fewer points as the $\lambda$ value increases because $\sum_1^4 \lambda_i = 1.0$.

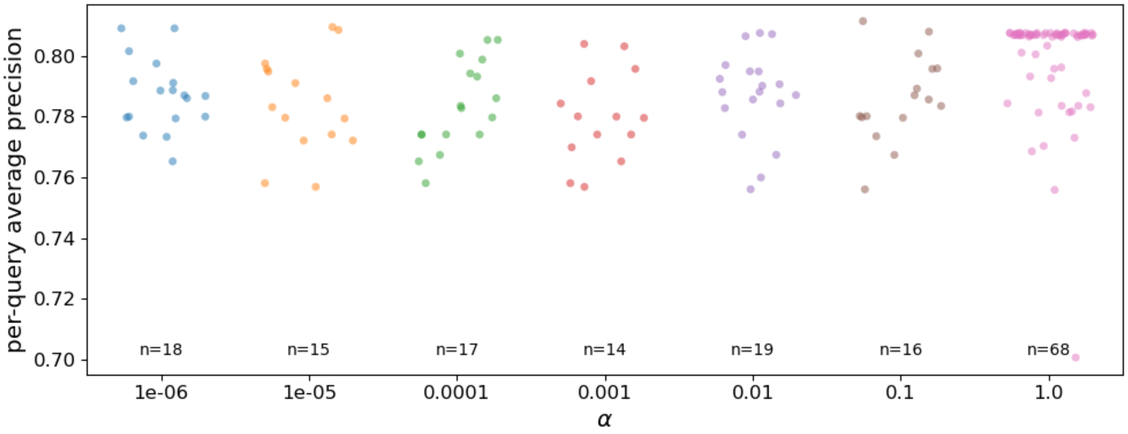


A

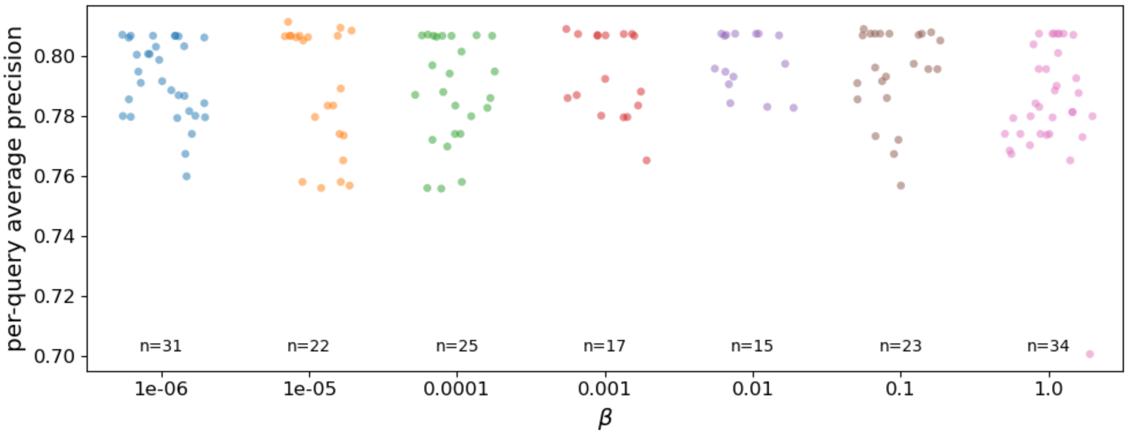


B

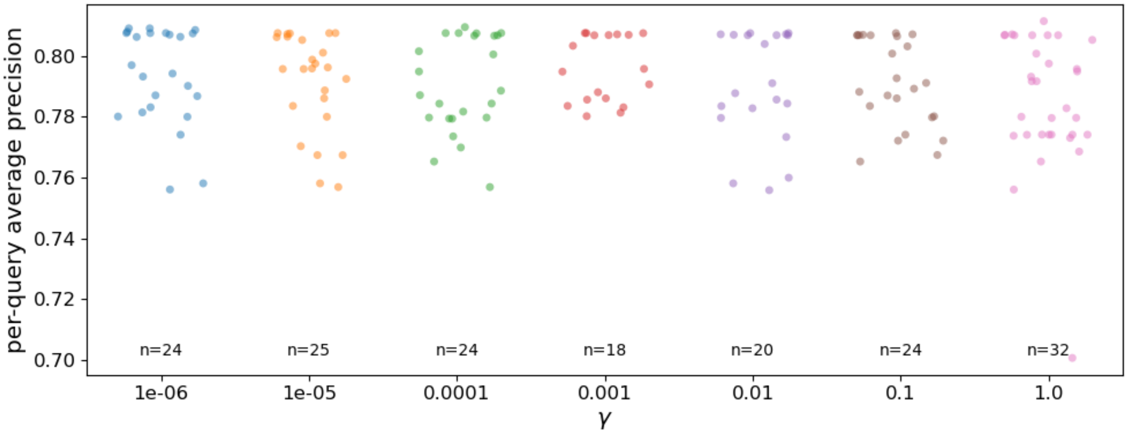


C

Supplemental Figure 10: **Performance as a function of $\alpha$, $\beta$, or $\gamma$.** This figures plots the performance of MS1Connect, given $N = 4000$ and $\delta_1 = 4$ ppm, as a function of (A) $\alpha$, (B) $\beta$, (C) or $\gamma$. In general, the values of $\alpha$, $\beta$, and $\gamma$ are not correlated with performance. Note that the values near the bottom are the number of points found in each column.

[4] H. Lin and J. A. Bilmes. Optimal selection of limited vocabulary speech corpora. In *Proc. Annual Conference of the International Speech Communication Association (INTERSPEECH)*, Florence, Italy, August 2011.

[12] The UniProt Consortium. UniProt: a worldwide hub for protein knowledge. *Nucleic Acids Research*, pages D506–D515, 2019.

[13] S. Pfrunder, J. Grossmann, P. Hunziker, R. Brunisholz, M. Gekenidis, and D. Drissner. Bacillus cereus group-type strain-specific diagnostic peptides. *Journal of Proteome Research*, 15(9):3098—3107, September 2016.
